# scTIE: data integration and inference of gene regulation using single-cell temporal multimodal data

**DOI:** 10.1101/2023.05.18.541381

**Authors:** Yingxin Lin, Tung-Yu Wu, Xi Chen, Sheng Wan, Brian Chao, Jingxue Xin, Jean Y.H. Yang, Wing H. Wong, Y. X. Rachel Wang

**Affiliations:** School of Mathematics and Statistics, The University of Sydney, NSW, Australia; Charles Perkins Centre, The University of Sydney, NSW, Australia; Laboratory of Data Discovery for Health Limited (D24H), Science Park, Hong Kong SAR, China; Department of Statistics, Stanford University, CA, USA; Institute of Electronics, National Yang Ming Chiao Tung University, Hsinchu, Taiwan; Department of Electrical Engineering, Stanford University, CA, USA; Department of Biomedical Data Science, Stanford University, CA, USA; Bio-X Program, Stanford University, CA, USA

**Author notes:** Equal contribution (Yingxin Lin, Tung-Yu Wu, Xi Chen). To whom correspondence should be addressed. Email: Y. X. Rachel Wang,; Wing H. Wong,.

**Keywords:** Single cell multiome, Temporal data integration, Context-specific gene regulatory network

## Abstract

Single-cell technologies offer unprecedented opportunities to dissect gene regulatory mechanisms in context-specific ways. Although there are computational methods for extracting gene regulatory relationships from scRNA-seq and scATAC-seq data, the data integration problem, essential for accurate cell type identification, has been mostly treated as a standalone challenge. Here we present scTIE, a unified method that integrates temporal multimodal data and infers regulatory relationships predictive of cellular state changes. scTIE uses an autoencoder to embed cells from all time points into a common space using iterative optimal transport, followed by extracting interpretable information to predict cell trajectories. Using a variety of synthetic and real temporal multimodal datasets, we demonstrate scTIE achieves effective data integration while preserving more biological signals than existing methods, particularly in the presence of batch effects and noise. Furthermore, on the exemplar multiome dataset we generated from differentiating mouse embryonic stem cells over time, we demonstrate scTIE captures regulatory elements highly predictive of cell transition probabilities, providing new potentials to understand the regulatory landscape driving developmental processes.

## Introduction

In eukaryotic cells, gene expressions are intricately regulated through complex interactions of transcription factors (TFs), various regulatory elements and target genes. Deciphering the functions of gene regulatory networks (GRNs) in shaping cell identity and cell fate is one of the central quests in understanding the mapping from genomic blueprints to phenotypes. Over the past decades, much effort has been devoted to developing statistical and computational methods for inferring GRNs from tissue-level bulk data containing genome-wide profiling of gene expression, TF binding, and 3D chromatin structure. More recently, the advent of single-cell sequencing technologies has propelled the study of GRNs into a new era, in which context-specific regulation mechanisms can be investigated. Such GRNs describe gene regulatory interactions that occur in a specific biological context, which may encompass different cell types, lineages, tissues, or environmental conditions. Alongside new opportunities, the sparse and noisy nature of these single-cell data also brings new challenges to the statistical and computational analyses.

A growing number of methods have been developed to extract GRNs from data generated by assays of single-cell RNA-sequencing (scRNA-seq) and single-cell transposase-accessible chromatin sequencing (scATAC-seq). Most of these methods infer the relationships between TFs and target genes by estimating their interactions with *cis*-regulatory elements (CREs) as an intermediate, using information including TF motif enrichment, marginal or conditional correlations between genes and CRE accessibility, and physical proximity between different elements [1, 2, 3, 4, 5]. These methods typically work with multimodal data that provide joint profiling of scRNA-seq and scATAC-seq from the same cells, or unpaired data from a matched population of cells, possibly measured over a time course. However, they do not directly address the data integration problem accompanying such data, in which noise, sparsity, and batch effects can obscure identification of cell types and affect the downstream inference of context-specific GRNs. Furthermore, to compare how GRNs dynamically evolve in developmental data, features (e.g., genes, CREs) that are different between time points (or pseudotime points) are identified using differential expression (DE) / accessibility (DA) analyses. While this captures marginal correlations, the features found are not necessarily predictive of the developmental changes.

On a separate front, an increasing number of computational methods have been proposed to perform data integration for single-cell multiomics data from unpaired measurements [6, 7, 8, 9]. As more technologies capable of multimodal profiling start to emerge [10, 11, 12], integration methods designed for paired data [13, 14, 15, 16] have also attracted significant research interests. However, most of these integration methods do not directly address the immediate downstream problem of inferring GRNs; one exception is GLUE [6], although the GRNs inferred there remain global and not context-specific. One difficulty lies in the fact that most of these methods rely on finding a low-dimensional representation of the datasets across modalities and data batches, and how to extract interpretable biological signals from blackbox methods such as neural networks is a challenging problem. Neural networks offer a conceptual advantage over methods built on linear models, including cross correlation analysis and non-negative matrix factorization, as their superior representation power can capture complex nonlinear interactions in the feature space. However, this comes with the drawback that the relationships between the measured features (e.g., genes) and cellular phenotypes in trained models become more difficult to interpret. Although alternative architectures have been proposed involving linearizing part of the neural network [17], a tradeoff remains between the network’s representation power and interpretability.

Here, we propose scTIE, an autoencoder-based method for integrating multimodal profiling of scRNA-seq and scATAC-seq data over a time course and inferring context-specific GRNs. To the best of our knowledge, scTIE provides the first unified framework for the integration of temporal data and the inference of context-specific GRNs that predict cell fates. We achieve this through three main innovations in the architecture design of the autoencoder and the interpretation of a blackbox neural network method. Firstly, scTIE uses iterative optimal transport (OT) fitting to align cells in similar states between different time points and estimate their transition probabilities. scTIE incorporates OT into the loss function of the autoencoder so that the alignment of cells is updated iteratively throughout training to achieve a desirable balance between time point alignment and cell type separation. This is in contrast to many widely used applications of OT in trajectory inference of scRNA-seq data [18, 19], where most of the methods solve OT only once on suitably constructed cell distance matrices. Secondly, scTIE removes the need for selecting highly variable genes (HVGs) as input through a pair of coupled batchnorm layers to account for large variations in gene expression levels, making it more robust and generalizable. Thirdly, scTIE provides the means to extract interpretable features from the common embedding space by linking the developmental trajectories of cell representations to their measured features (genes and peaks). We formulate a trajectory prediction problem using the estimated transition probabilities from OT and use gradient-based saliency mapping [20, 21] to identify genes and peaks that are potentially driving the cellular state changes.

To demonstrate the performance of scTIE on developmental data, we have chosen to focus on multimodal time-course data, as this emerging form of data provides better opportunities to understand the key transcriptional regulatory activities driving a developmental process. To assess scTIE’s integration performance against other existing methods, we constructed a variety of synthetic datasets using a mouse early organogenesis multiome dataset. We show that scTIE effectively aligns cells from different time points and removes batch effect, providing an optimal tradeoff between time alignment, modality alignment and cell type separation. We further generated an exemplar dataset comprising paired scRNA-seq and scATAC-seq measurements from *∼* 11, 000 differentiating mouse embryonic stem cells (mESCs) over a time course. Applying scTIE, we show its superior capacity to capture biological signals from each modality and achieve better day alignment when compared to other methods, resulting in identification of distinct cell subpopulations. Finally, using developmental transitions from anterior primitive streak as a case study, we demonstrate scTIE’s ability to construct lineage-specific GRNs consisting of regulatory elements with a high predictive power of cell fate and identify key regulatory signals that would be missed by DE or DA-based analysis.

## Results

### Overview of scTIE

scTIE uses modality-specific encoders and decoders to project high dimensional input data from all time points into a lower dimensional common embedding space and reconstruct them in the original space (Fig. 1). A modality alignment loss is used to ensure the projected feature vectors from the same cell are close in distance. Each encoder-decoder pair is designed to preserve the original dimension of the input data with minimal information loss. For scATAC-seq, accessibility peaks are used as input without conversion to gene activity scores. The encoder and decoder for scRNA-seq use an additional pair of coupled batchnorm layers to handle heterogeneity in gene expression levels and achieve high-fidelity reconstruction of the signals without the need for selecting HVGs. Between consecutive time points, scTIE models cell trajectories using the principle of OT based on the current embeddings and computes an OT loss using the transport cost matrix. The OT loss is incorporated into the total loss function to update the embedded features, aligning cells by their estimated transition probabilities in the trajectories; the cost matrix itself is also updated iteratively throughout training. Finally, scTIE finetunes the learned embeddings to build a supervised model for predicting cellular transition probabilities for subgroups of cells. Genes and peak regions highly predictive of the cellular transitions are selected by backpropagating the gradients, allowing us to construct GRNs responsible for developmental changes.

**Figure 1:**
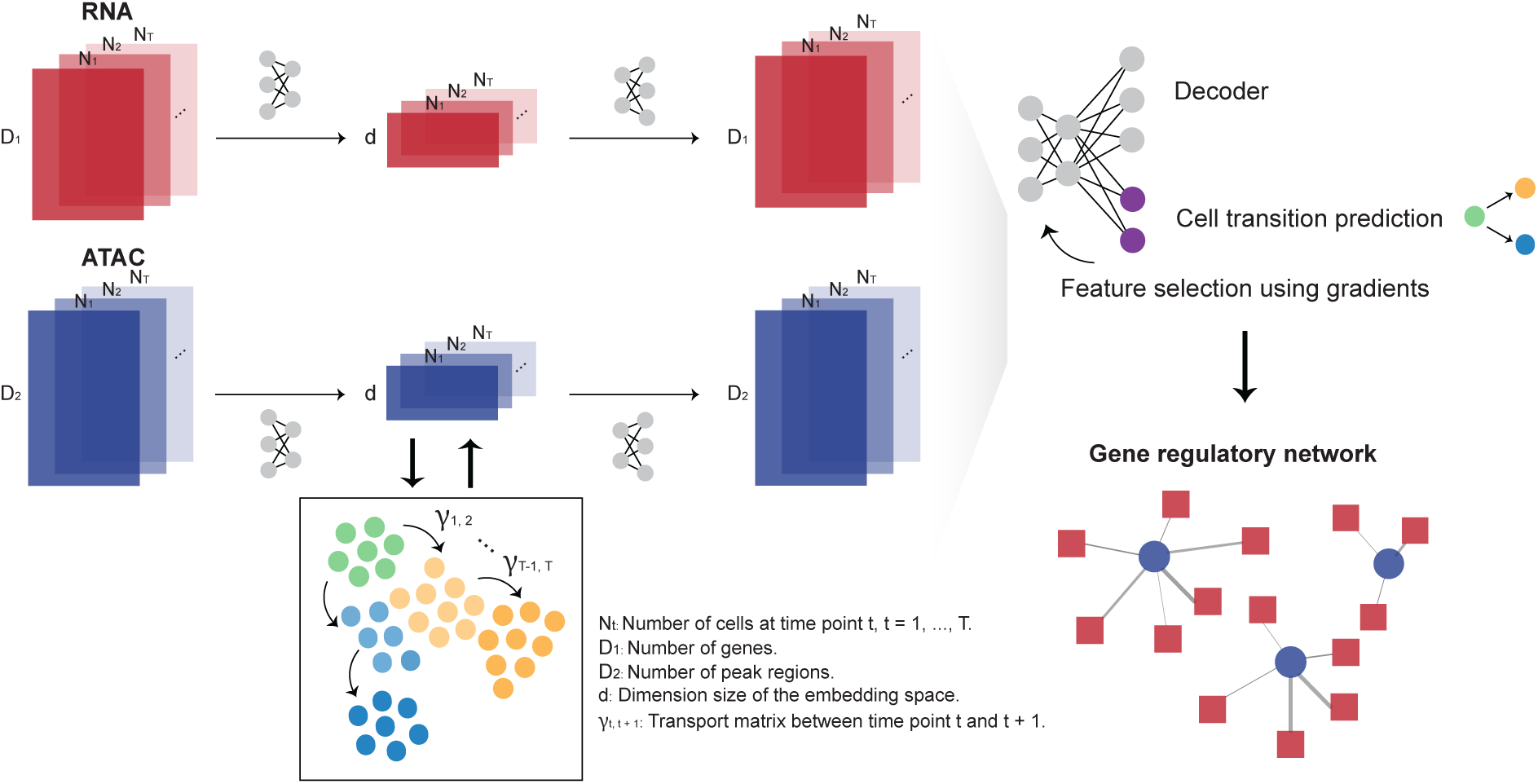
Overview of scTIE, a unified framework for the integration of temporal data and the inference of context-specific GRNs that predict cell fates. The input of scTIE consists of the gene expression matrix of scRNA-seq and peak matrix of scATAC-seq from single-cell multiome data over a time course.

### scTIE outperforms existing methods in integrating temporal multimodal data

We first evaluated the data integration performance of scTIE against recent methods designed to integrate paired multimodal data, including Seurat [15], scAI [16], multiVI [14] and MOFA [13]. We generated four synthetic datasets by introducing batch effects and noise into a mouse early organogenesis multiome dataset [22] (Fig. 2A, Supplementary Fig. S3). As shown in the UMAP plots of the data with synthetic batch effects introduced in RNA and noise introduced in ATAC (Fig. 2A), scTIE effectively removed the batch effects while also better revealing the cell type signals.

**Figure 2:**
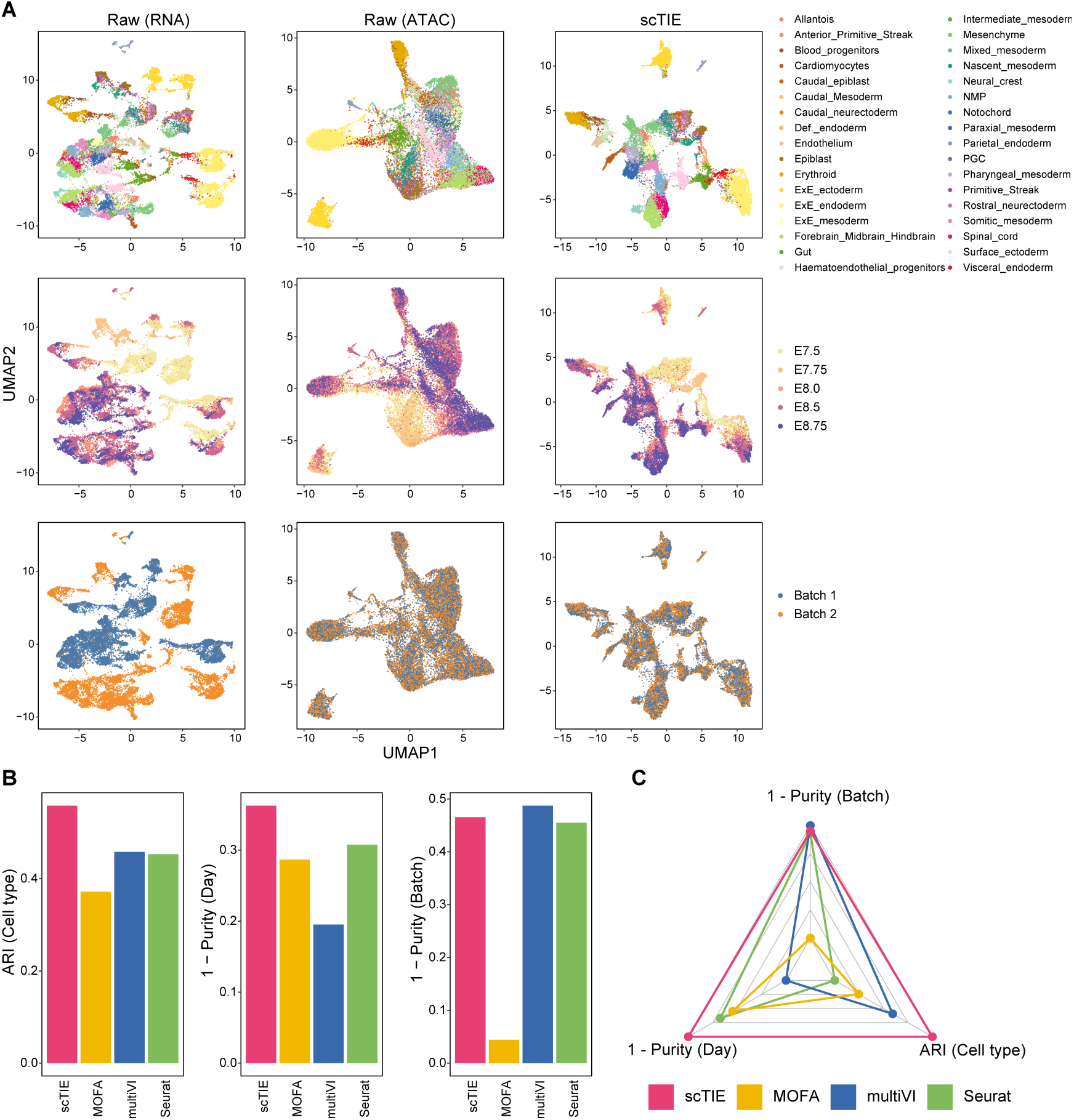
(A) Joint visualization using UMAP of the synthetic dataset with batch effect in RNA and noise in ATAC, colored by cell type annotations (first row), sampling days (second row) and synthetic batch information (third row). Each dot represents a cell in the embedding space. (B) Bar plots showing the evaluation metrics of different data integration methods, including ARI values for clustering with annotations (left); 1 – average purity scores of sampling days with the number of neighbors equal to 50 (middle) and 1 – average purity scores of the synthetic batch with the number of neighbors equal to 50 (right). Higher values indicate better agreement with annotations and mixing of batches/days. (C) Radar plot summarizing the three evaluation metrics shown in (B), where each line represents the performance of one method, and each axis represents an evaluation metric, starting from the minimum value of all methods. It is noted that scAI was not included in this benchmarking due to its long computational time (*>* 2 days).

Next, we compared the performance of these methods from three aspects, namely batch effect removal, time point alignment and their ability to capture cell type signals. We quantify the quality of batch removal and time point alignment using purity scores, which calculate the proportion of cells from the same batch/sampling time among neighbors of given cells. A lower purity score indicates a better mixing of batch/time points. We measured the cell type preservation using adjusted rand index (ARI) with the cell type annotations provided in the original paper as the ground truth. We find that scTIE outperforms the other methods in the overall performance across the three metrics (Fig. 2 B-C and Supplementary Fig. S1). Furthermore, scTIE’s superior performance is robust against the number of neighbors used in the purity score calculation (Supplementary Fig. S2). We observe similar trends across the other three synthetic scenarios, where scTIE consistently exhibits better performance than the other methods (Supplementary Fig. S3). Together, we demonstrate the superiority of scTIE in data integration, enabling better capture of biological signals through batch effect removal and time point alignment.

### scTIE enables identification of cellular subpopulations via modality and time point alignment with robust performance

Encouraged by scTIE’s performance in data integration, we next generated a temporal single-cell multimodal dataset and leveraged scTIE for the integration of cells across time points and annotation of cell types. We performed single-cell multiome sequencing from mESCs treated with Activin A/Lithium Chloride and measured on Day 2, 4 and 6, using the 10x Chromium Single Cell Multiome platform. After quality control filtering (Supplementary Fig. S4), we obtained high quality measurements of RNA and ATAC from a total of 11,440 cells, with a median detection of 4,130 genes expressed per cell and a median of 11,267 peaks detected per cell.

By clustering on the joint embeddings produced by scTIE, we identified 17 clusters with either distinct transcription or chromatin accessibility profiles that include cell types from all the three germ layers as well as from extra-embryonic layers of embryonic development (Fig. 3A-C). We annotated these clusters based on the key markers identified in the two previous studies [23, 24] (Fig. 3C), and confirmed them by label transfer using a public reference [25, 23] (Supplementary Fig. S5). Further explorations of the motif enrichment of regions with DA in specific clusters highlight the cluster-specific TFs of the annotated cell types (Fig. 3D-E). Additionally, we quantitatively assessed the clustering results using evaluation metrics. Our findings demonstrate that scTIE better preserves biological signals in each modality and achieves better alignment in days compared with the existing methods, further supporting our annotation of the cells using the integrated data from scTIE (Supplementary Figs. S8-S9).

**Figure 3:**
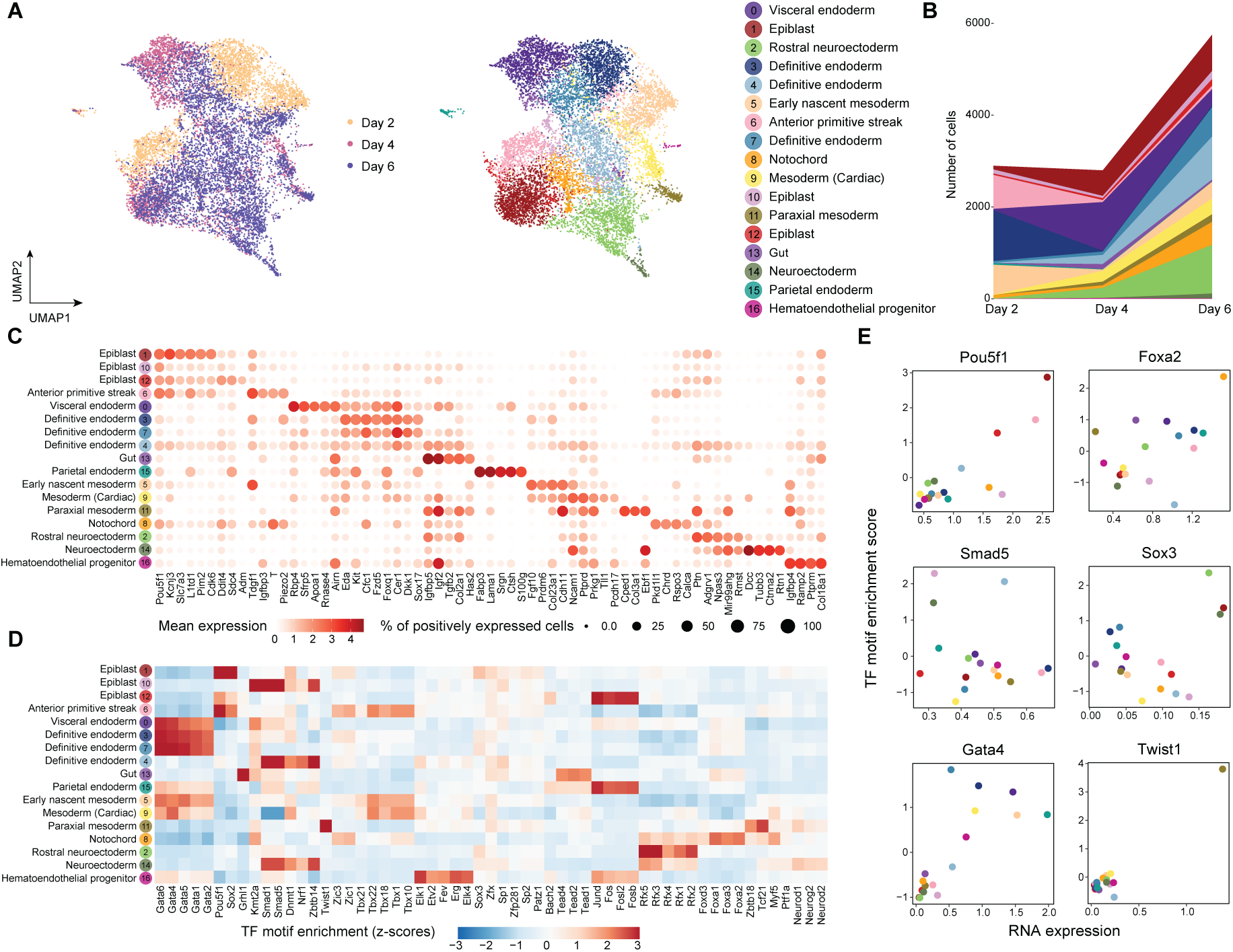
(A) Joint visualization of ESC dataset using UMAP, colored by sampling day and cell type annotations. Each dot represents a cell in the embedding space. (B) Cell type compositions per time point. (C) Dot plots of mean expression of RNA data. Rows represent cell types and columns indicate each genes. The color scale represents the expression level, and the size indicates proportion of positively expressed cells. The five most significantly expressed genes for each cluster are included. (D) Heatmap of the TF motif enrichment (z-scores) of ATAC data. Rows represent cell types and columns indicate TFs. The five most significantly enriched TFs for each cluster are included. (E) Scatter plots of the mean RNA expression levels by clusters (x-axis) and the average TF motif enrichment scores of ATAC (y-axis) for the selected TFs. The dots are colored by the cell type annotations, with color legend consistent with Fig. 3A.

Notably, scTIE identifies three distinct clusters of definitive endoderm (Cluster 3, 4 and 7) (Supplementary Fig. S6A). We find that Cluster 4 uniquely expresses several Wnt pathway direct targets (Vcan, Nrcam and Ccnd2) and Wnt TF (Lef1), and has lower expressions in Wnt inhibitors Dkk1 and some definitive endoderm markers (Hhex and Sox17) (Supplementary Fig. S6B). The activation of Wnt signaling of this group of cells could be linked to primordial lung specification progenitors [26]. Cluster 3 and Cluster 7 have similar expression profiles to each other. Compared with Cluster 3, we find Cluster 7 with majority of cells from Day 6 has lower expressions in Nodal signaling genes Nodal and Tdgf1, but higher expressions in genes that negatively regulate the Nodal pathway (Cer1 and Lefty1) (Supplementary Fig. S6B).

An inspection of the epiblast subsets further demonstrates that scTIE enables cellular subpopulation identification (Supplementary Fig. S7A). We find that one of the epiblast clusters (Cluster 12) has upregulation of genes related to Hypoxia (Adm, Anxa2, Ddit4 and Gbe1), which could enhance the defintive endoderm differentiation, as suggested in [27, 28] (Supplementary Fig. S7B). In addition, we find that Cluster 1 is enriched with anterior epiblast markers (Pou3f1, Enpp3, Pten and Slc7a3), while Cluster 10 highly expresses posterior epiblast markers (Lhx1, Ifitm1) (Supplementary Fig. S7B) [29], with downregulation of the TFs Pou5f1 and Sox2 but upregulation of the TFs Foxa1 and Foxa2 (Supplementary Fig. S7C).

Finally, we examine the stability of our results in both modality alignment and cluster identification, with respect to key tuning parameters in scTIE, including the weight of OT in the loss function, the number of nodes in hidden layer and the updating frequency of OT. We find that the weight of the OT loss is an important parameter to reach a balance between the alignment of modalities and time points, with a larger weight resulting in a better alignment in time points (Supplementary Fig. S11A) but poorer performance in modality integration (Supplementary Fig. S10A, D). In this sense, the choice of this parameter can be guided by the performance in modality alignment, since the pairing information for all cells is known and serves as the ground truth. The two other tuning parameters have a small impact on our results (Supplementary Fig. S10BC, E-F, Supplementary Fig. S11B-C).

Together, we demonstrate that scTIE is able to capture distinct cellular subpopulations by preserving information from both epigenomic and transcriptomic profiles, while also aligning the cells from different time points.

### scTIE embeddings capture interpretable biological features

To interpret the embedding space projected by scTIE, we deconvoluted the latent representation by backpropagating the gradient of each dimension in the embedding layer with respect to gene and peak input, followed by ranking the features. We then computed the enrichment scores of the cell type marker list for the feature rankings of each embedding dimension (see Methods). We find that each dimension exhibits distinct patterns of enrichment of cell type markers, and at the same time the cell types from the same lineage share similar enrichment patterns across the dimensions, indicating that scTIE captures diverse and biologically meaningful information from the data (Fig. 4A). We further observe that the enrichment results of RNA and ATAC share similar patterns, illustrating that scTIE is able to link the transcriptomic profiles with the chromatin accessibility through the common embeddings (Fig. 4A).

**Figure 4:**
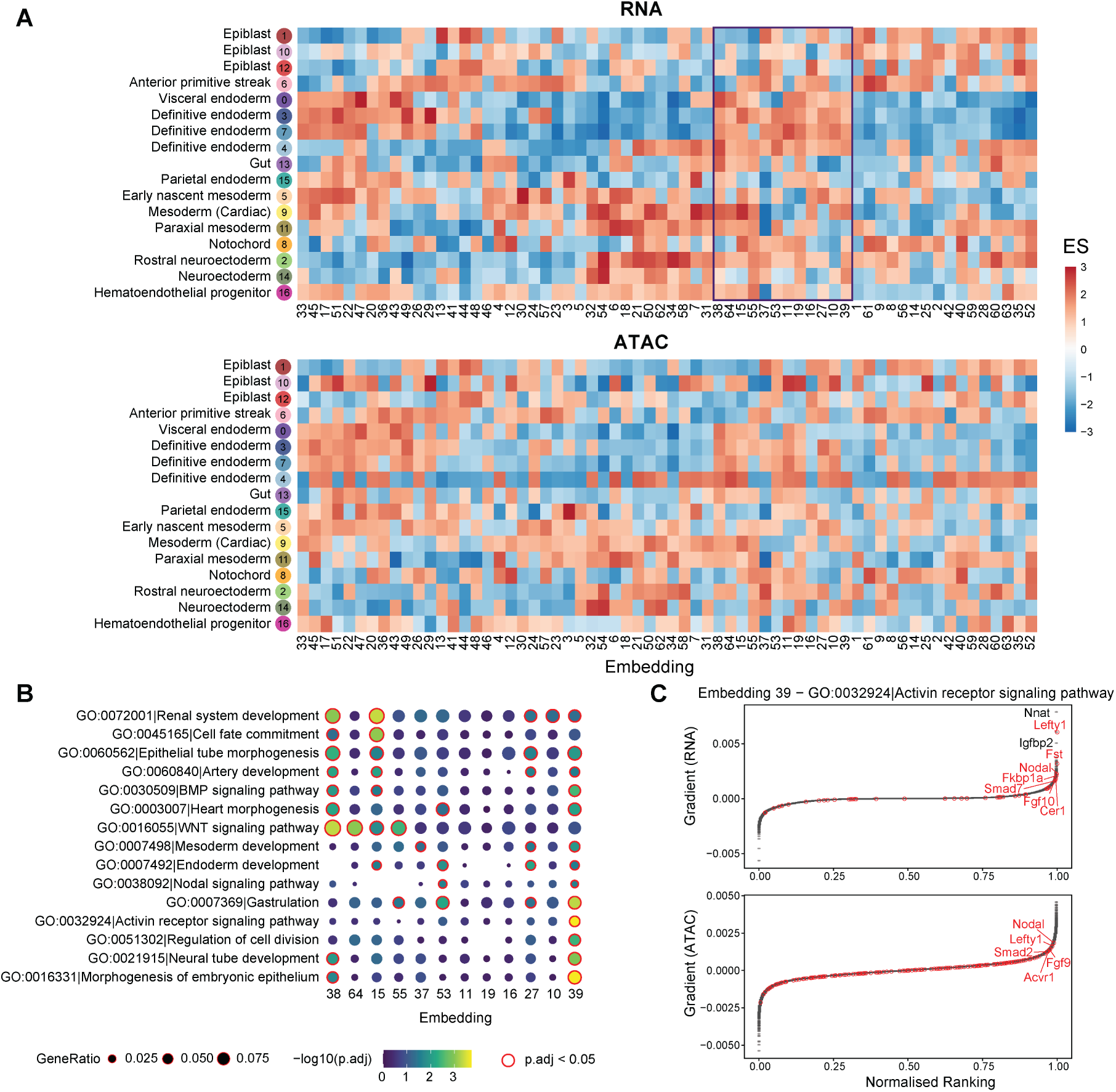
(A) Enrichment scores of the gradient ranking in each embedding dimension using the RNA (top panel) and ATAC (bottom panel) marker list for each cell type. (B) Gene ontology enrichment of selected pathways on the gradient ranking of a subset of embedding dimensions. (C) Gradient rankings for RNA (top panel) and ATAC (bottom panel) of embedding dimension 39, where genes/peaks are ranked based on the gradient values. The labeled points are genes in the selected gene set (Activin receptor signaling pathway).

The embedding gradients can be further interpreted in terms of known biological functions, based on their Gene ontology (GO) enrichment. As illustrated in Fig. 4B, we find that the embedding dimensions enriched with definitive endoderm cell type markers can be associated with different pathways. Interestingly, we observe that dimension 39 is uniquely enriched with Activin receptor signaling, as confirmed by the top ranking genes including Lefty1, Fst, and Nodal from this pathway (Fig. 4C). Consistently, the nearest genes of the top ranking peaks also include genes associated with the Activin pathway, such as Nodal, Lefty1 and Fgf9. Since treatment by Actinvin is a key component of our differentiation protocol (see Methods), it is comforting to see that the relevance of this pathway is captured by the fitted model. Together, we demonstrate that scTIE is able to project the two modalities into a joint embedding space that captures interpretable biological signals of the data.

### scTIE uncovers cell fate-specific regulatory networks

scTIE constructs lineage-defining GRNs by combining information across different dimensions of the embedding layer to predict the cell transition probabilities between time points. As a case study, we investigate the transitions of cells from anterior primitive streak on earlier days into endoderm, mesoderm, as well as remaining as anterior primitive streak on later days. The primitive streak is a transient embryonic structure which marks bilateral symmetry, helps confer anterior-posterior spatial information during gastrulation, and initiates germ layer formation [30]. A distinct group of cells located at anterior primitive streak, the node, forms the axial mesodermal structures and definitive endoderm cells [31].

In each of the above three possible cell fates, we fine-tuned the trained embeddings using a prediction layer with weight regularization and backpropagate the gradients from the prediction layer to select the top 200 genes and 500 peak regions as the most predictive features of the lineage. Compared with the conventional approach that uses DE / DA analysis to select the top features, scTIE selects genes and peak regions with significantly better prediction performance (Fig. 5A). The superior prediction performance is consistent across a range of tuning parameters, including the regularization weights and the number of top features, evaluated via cross validation (Supplementary Fig. S12).

**Figure 5:**
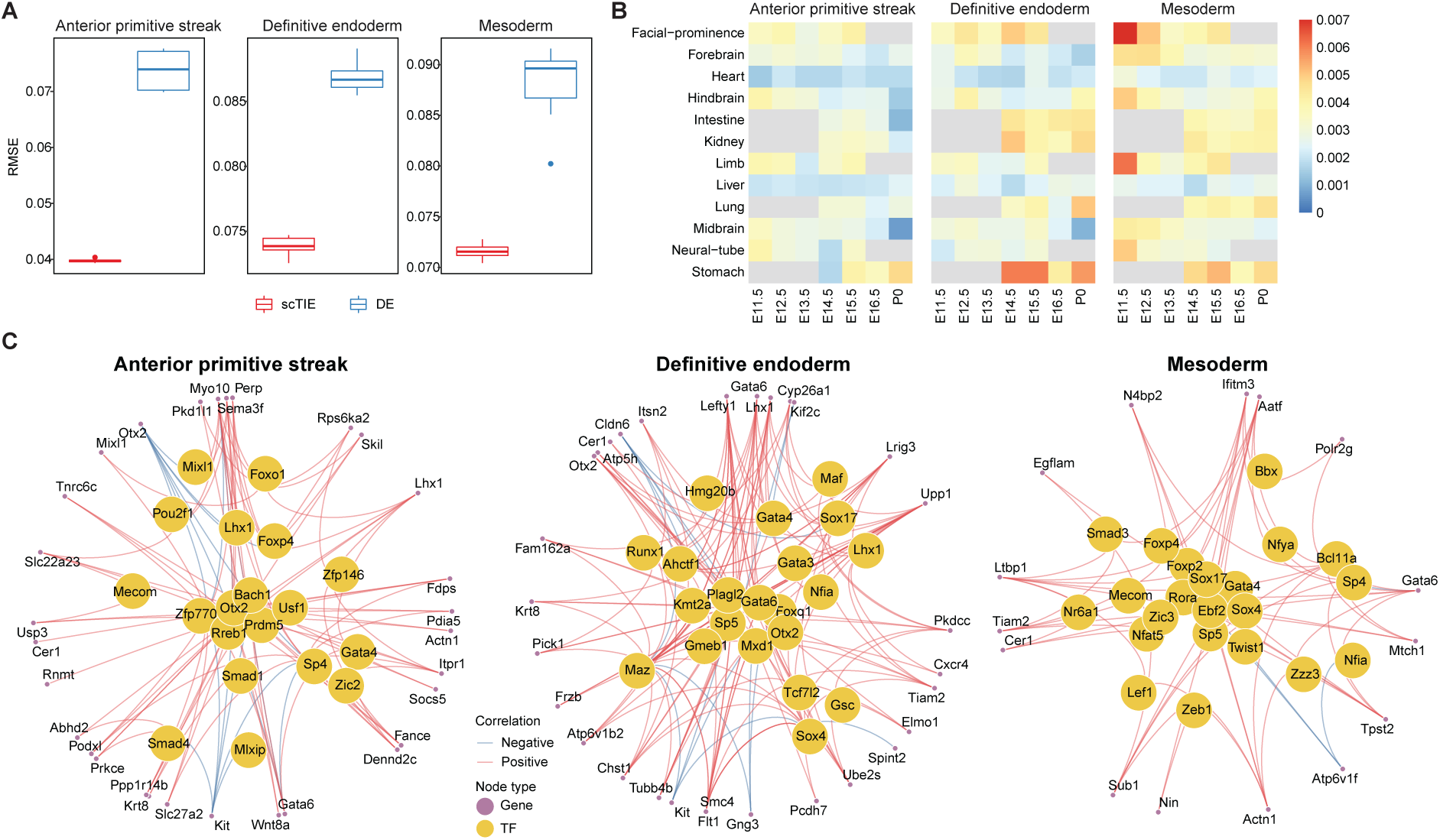
(A) Performance of cell fate probability prediction. (B) Similarity of top gradient peaks with enhancers of 12 tissues at seven developmental stages from known enhancer databases. (C) GRN of three cell fates.

To annotate the top peaks, we overlapped the selected peaks with the published enhancer database from 12 tissues of seven developmental stages from 11.5 days after conception until birth [32], quantified by the Jaccard index. We find that the top peaks associated with mesoderm transition potential are enriched with facial prominence and limb enhancers at E11.5, while endoderm transition-related peaks identified by scTIE show higher enrichment and distinct overlap with stomach enhancers at E14.5, E15.5 and P0 (Fig. 5B). In contrast, the peaks selected by DA analysis show enrichments in tissues that are much less specific to predicted lineages of mesoderm or endoderm (Supplementary Fig. S13). Together, these results illustrate that scTIE is able to identify peaks that are specific to lineage transition.

The identification of genes and peaks that are predictive of cell transition further allows us to infer GRN for each of the lineages: anterior primitive streak, endoderm and mesoderm (see Methods). In the GRN of anterior primitive streak (Fig. 5C, left panel), we identified a few TFs that play key roles in jointly governing anterior mesendoderm and the node development (Lhx1, Otx2 and Smad4) [33, 34], as well as a TF related to axial mesendoderm morphogenesis and patterning (Mixl1) [35]. Interestingly, when focusing on the endoderm GRN (Fig. 5C, middle panel), we find that besides identifying TFs that are central regulators for the formation of definitive endoderm development (Sox17, Gata4, Gata6, and Gsc) [36, 37, 38, 39, 40], scTIE also captures TFs that are associated with early mesendoderm differentiation (Runx1) [41] and morphogenetic movement (Lhx1) [42].

Lastly, we examined the mesoderm GRN (Fig. 5C, right panel) which identifies a few key TFs (Hhex, Sox17, Smad3, Zic3, Twist1 and Nfat5) that are associated with mesoderm lineages. Notably, most of these TFs have insignificant p-values under DE analysis (Table S1), illustrating that scTIE captures key regulatory signals in this lineage that would be missed otherwise. More specifically, the mesoderm GRN highlights TFs that are associated with cardiac development such as Zic3 in early mesodermal patterning [43, 44]; Hhex that is involved in mediating the Sox17 for cardiac mesoderm formation in mESC [45] and Nfat5 for cardiomyogenic during mesodermal induction through regulating the canonical Wnt pathway [46]. We also identify TFs that are essential for mesoderm formation and patterning (Smad3) [47] and cranial mesoderm development (Twist1) [48].

## Discussion

While the rapidly increasing collection of single-cell multiomics data provides a wealth of information for examining context-specific regulatory mechanisms, accurate characterization of cell identities remains the first hurdle to be overcome in such tasks. scTIE provides a unified framework for the integration and joint modeling of temporal multimodal data and the subsequent visualization, cell type identification and inference of key regulatory modules predictive of the developmental transitions of cells. Incorporating OT into the training of an autoencoder, scTIE alternates between updating the alignment of cells at different time points and using the current alignment for training the projections into the common embedding space, thus achieving a better balance between integrating time points and maintaining cell type specific signals. As we have demonstrated on the real and synthetic datasets, scTIE outperforms existing paired methods in terms of integration performance.

Different from existing integration methods that also utilize the notion of a common embedding space, scTIE directly exploits the information in this space produced by the nonlinear projections of a neural network, linking it to interpretable features such as genes and peak regions. scTIE extracts context-specific gene regulatory relationships through the identification of features that are predictive of cell transition probabilities, which quantify how likely a collection of cells on earlier days will transit to a certain cell state on later days, relative to other cells. These sets of cells can be flexibly defined, allowing users to investigate any cell transition process of interest. In addition to cell transition probabilities derived from OT, the current framework can also be adapted to select features that are predictive of other types of response variables, such as pseudotime and perturbation, which potentially enables the construction of differential GRN under continuous cell differentiation and in perturbed conditions.

scTIE is designed for temporal multimodal data, which is ideal for studying single-cell genomics in developmental trajectories. Paired measurements from the same cells remove the need for computational pairing, which can introduce errors into the downstream GRN analysis if cells of different cell types are paired, and the issue of cell type imbalance between different modalities. The integration of unpaired developmental data across multiple time points remains an open problem itself. For datasets taken from a matched population, a loss function performing global alignment between modalities, such as the one used in [9], can be potentially incorporated into the training of scTIE. However, the problem is more challenging if cells are sampled at different time points or develop at a different rate across the modalities, and we will pursue this in future work.

Although a large number of methods exist for inferring pseudotime ordering of cells from a static snapshot of a developmental process, pseudotime inference assumes that a continuum of cellular states is observed at the sampled time, and thus may not capture the entire transition process [49]. An interesting extension would be combining pseudotime inference and experimental time points to create a finer temporal resolution. However, we note that this would also increase the computation time of scTIE, since iterative OT estimation is performed between consecutive time points; efficient and accurate OT algorithms remain an active area of research.

We have focused on scRNA-seq and scATAC-seq as common modalities from multimodal profiling technologies. Other modalities such as methylation and protein levels [50, 51, 52] can be easily incorporated into scTIE through appropriate encoder-decoder pairs. Since transcriptional regulation involves interactions of protein complexes, histone modifications and other microenvironmental factors, we expect the addition of such information will allow us to build a more accurate prediction model for cellular state changes. Furthermore, emerging single-cell perturbation assays [53] can either be used to validate the top candidates found in our predictive model, or built into the neural network architecture as a prior knowledge graph [6].

In summary, scTIE provides an integrative framework for analyzing temporal multimodal data, which is an emerging form of data we expect will become more readily available as interests in characterizing GRNs at single-cell resolution continue to rise. On real and synthetic developmental datasets, scTIE is shown to provide effective integration of cells from all time points and select key regulatory elements with superior performance in predicting cellular state changes. We envision that advances in single-cell technologies generating new forms of temporal data will enable us to further expand the functionalities of scTIE, paving the way towards a holistic understanding of cellular transitions and responses in development and disease.

## Methods

### Synthetic data construction

The 10x Genomics multiome data of mouse early organogenesis, along with its cell type annotation, was obtained from the Gene Expression Omnibus database under accession number GSE205117 [22]. The dataset comprises 59,132 cells from a time course of mouse embryonic development, spanning 5 time points from E7.5 to E8.75.

To construct synthetic data that could be processed by most of the methods within their computational capacity, we subset the data to 24,188 cells by selecting only one sample at each time point. We filtered out genes expressed in less than 1% of cells and peaks expressed in less than 5% of cells, resulting in 15,754 genes and 81,108 peaks. To introduce noise and batch effects to the data, we used the `downsampleReads()` function in the DropletUtils R package to downsample the reads. We generated four synthetic scenarios: (1) subsample 10% for all cells in ATAC; (2) subsample 10% for all cells in ATAC and 50% for all cells in RNA; (3) subsample 50% for half of cells in RNA to create the synthetic batch effect in the data; and (4) subsample 10% for all cells in ATAC, subsample 50% for half of the cells in RNA and 25% for the other half of the cells.

### mESC data generation

#### Cell culture

Mouse embryonic stem cell line R1 was obtained from ATCC. The cells were first expanded on an MEF feeder layer previously irradiated. Then, subculturing was carried out on 0.1% bovine gelatin-coated tissue culture plates. The cells were propagated in mESC medium consisting of Knockout DMEM supplemented with 15% Knockout Serum Replacement, 100 *µ*M nonessential amino acids, 0.5 mM beta-mercaptoethanol, 2 mM GlutaMax, and 100 U/mL Penicillin-Streptomycin with the addition of 1,000 U/mL of LIF (ESGRO, Millipore).

#### Cell differentiation

mESCs were differentiated using the hanging drop method [54]. Trypsinized cells were suspended in chemically defined medium CDM [36] to a concentration of 37,500 cells/mL. CDM consists of 75% Iscove’s modified Dulbecco’s medium (IMDM, Invitrogen), 25% Ham’s F12 medium (Invitrogen), 1X N2 supplements (Invitrogen), 0.05% bovine serum albumin (BSA, Invitrogen), 2 mM Glutamax-1 (Invitrogen), 0.5 mM ascorbic acid (Sigma-Aldrich), and 4.5 x 10^4^ M MTG (Sigma-Aldrich). 20 *µ*L drops (*∼*750 cells per drop) were then placed on the lid of a bacterial plate and the lid was upside down. After 48 h incubation at 37*^◦^*C incubator with 5% CO_2_, Embryoid bodies (EBs) formed at the bottom of the drops were collected and placed in the well of a 6-well ultra-low attachment plate (Corning) with fresh CDM medium containing 50 ng/mL Activin A (R&D Systems, 338-AC-050/CF) and 2 mM Lithium Chloride (LiCl, Sigma-Aldrich) for up to 6 days, with the medium being changed daily.

#### Single cell multiome library

We followed 10x Genomics single cell multiome library preparation protocol. The EBs were collected at Day 2, 4, and 6 after Activin A/Lithium Chloride treatment. For each time point, the cells were first treated with StemPro Accutase Cell Dissociation Reagent (Thermo Fisher) at 37*^◦^*C for 10-15 min with pipetting. Single cell suspension was obtained by passing through 37 *µ*M cell strainer (STEMCELL Technologies) twice. After measuring cell concentration, approximately 1 million of cells were centrifuged at 300 rcf for 5 min. Nuclei were isolated by following the protocol provided by 10x Genomics (Nuclei isolation for single cell multiome ATAC + Gene expression sequencing, CG00365, Rev A). The final nuclei concentration was adjusted to 3000 cell/*µ*L in 1X Nuclei Buffer (10x Genomics). The sample was immediately submitted to Stanford Genomics Service Center (SGSC) for single cell sorting using 10x Chromium Controller (target cells: 5000 per replicate, total 2-3 replicates per time point). The singe cell multiome library was generated using Chromium Next GEM Single Cell Multiome ATAC + Gene Expression Reagent Bundle Kit (10x Genomics, PN-1000283).

#### Data preprocessing

10x Genomics Cell Ranger arc v2.0.0 was used to process the raw fastq files for each multiome single-cell dataset separately. The reference genome and transcriptome for alignment and annotation was version arc-mm10-2020-A-2.0.0. To integrate all filtered count matrices for scRNA-seq and scATAC-seq from different replicates and time points, the cellranger-arc aggr command was applied with default depth normalization method.

Next, we performed quality control on the cell level. We removed cells based on the following criteria in scRNA-seq: (1) with the total number of UMI (nUMI) less than 6000 on Day 2, 3000 on Day 4 and Day 6; (2) with nUMI greater than 100,000; (3) with the number of genes less than 2000 on Day 2, 1800 on Day 4 and 1500 on Day 6 and (4) mitochondrial reads greater than 25%. We further removed cells based on the following criteria in scATAC-seq: (1) with less than 500 total ATAC fragments and (2) with less than 500 peaks detected. After quality control, we retained 11440 cells (Day 2: 2896 cells; Day 4: 2796 cells and Day 6: 5748 cells). We then performed the quality control on the feature level, removing the genes that are not expressed in any cells and the peaks that are expressed at least 5% of cells, resulting in 26717 genes and 61744 peaks as input in scTIE.

### Architecture and training of scTIE

scTIE uses an autoencoder structure to project high dimensional feature vectors (i.e., gene expression levels and accessibility peaks) from all time points into a lower dimensional common embedding space and reconstruct the features in the original high dimensional space. Each modality has its own encoder and decoder (Table 1). For RNA, the architecture has an additional pair of coupled batchnorm layers, where the final reconstructed output uses the moving average *µ* and standard deviation *σ* stored in the first batchnorm layer of the encoder to perform rescaling. This accounts for the high variability in gene expression levels without the need for selecting HVGs, and allows us to significantly improve the performance in reconstruction correlation, modality and day alignment, and clustering quality (Supplementary Fig. S14). The pairing between feature vectors from the same cell is enforced through a modality loss function minimizing their distance in the embedding space. An OT matrix is used to construct cell trajectories between each pair of consecutive time points. In contrast to existing methods using OT for trajectory inference, we integrate an OT loss into the autoencoder training process and estimate the OT matrix iteratively throughout. A larger weight on the OT loss leads to better alignment between days (Supplementary Fig. S11A).

**Table 1:**
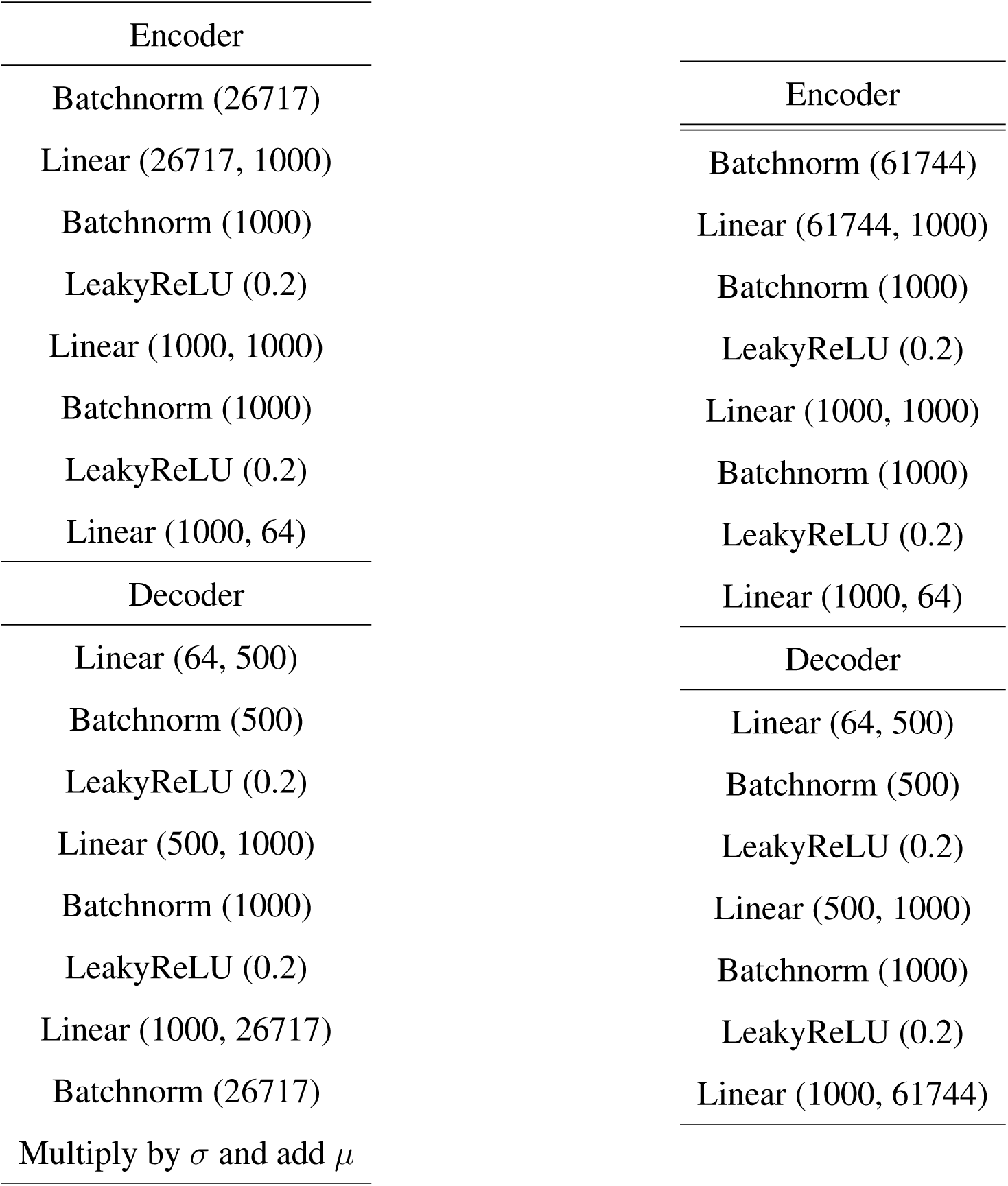
Autoencoder architecture for RNA (left) and ATAC (right).

| Encoder | Encoder |
| --- | --- |
| Batchnorm (26717) | Batchnorm (61744) |
| Linear (26717, 1000) | Linear (61744, 1000) |
| Batchnorm (1000) | Batchnorm (1000) |
| LeakyReLU (0.2) | LeakyReLU (0.2) |
| Linear (1000, 1000) | Linear (1000, 1000) |
| Batchnorm (1000) | Batchnorm (1000) |
| LeakyReLU (0.2) | LeakyReLU (0.2) |
| Linear (1000, 64) | Linear (1000, 64) |
| Decoder | Decoder |
| Linear (64, 500) | Linear (64, 500) |
| Batchnorm (500) | Batchnorm (500) |
| LeakyReLU (0.2) | LeakyReLU (0.2) |
| Linear (500, 1000) | Linear (500, 1000) |
| Batchnorm (1000) | Batchnorm (1000) |
| LeakyReLU (0.2) | LeakyReLU (0.2) |
| Linear (1000, 26717) | Linear (1000, 61744) |
| Batchnorm (26717) |  |
| Multiply by $\sigma$ and add $\mu$ | |

Let *X*^(*t,s*)^ denote the data matrix from time point *t* and modality *s*, where *t* = 1*, …, T* and *s* = 1, 2 for RNA and ATAC respectively. Each time point *t* provides measurements for *N_t_*cells; thus in this case, 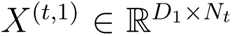 with *D*_1_ = number of genes and 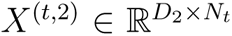 with *D*_2_ = number of peak regions. In each iteration, a mini-batch of data is sampled by taking equal-sized subsets of cells from each time point, that is, 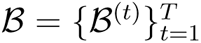, where each subset *B*^(*t*)^ has *B* cells. Three loss functions are applied to the mini-batch.

1. *Reconstruction loss*. (*f_s_, g_s_*) represents the encoder-decoder pair for modality *s*. Compared with the architecture for ATAC, the RNA part has a pair of coupled batchnorm layers, starting with a batchnorm layer in the encoder to remove scale variations in genes and prevent the gradients from being dominated by a small number of highly expressed genes (Table 1). Let 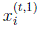 denote the gene expression vector from cell *i* at time *t* and 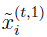 denote the normalized output from the first batchnorm layer, then 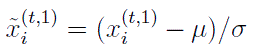 where *µ* and *σ* are the moving average and standard deviation of the genes saved in the batchnorm layer throughout training. The reconstruction loss is applied to the normalized data and the output from the decoder, defined as

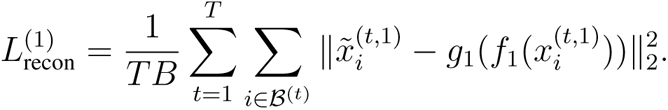

For ATAC, the first layer in the encoder is a fully connected layer and the reconstruction loss is computed on the input 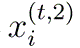 and output 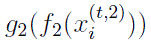 as usual. The overall *L*_recon_ is the sum of 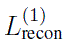 and 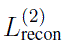.
2. *Optimal transport loss*. We leverage OT to effectively align cells from all time points in the embedding space. For notational convenience, we will suppress the dependence on modality *s* for now, with understanding that the following steps are performed for each modality. For any two adjacent time points *t* and *t* + 1, a transport cost matrix 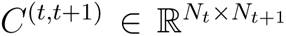 can be computed using the current embeddings, where the (*k, l*)-th entry is given by 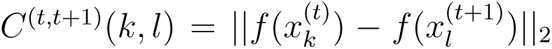 for the *k*-th cell from *t* and the *l*-th cell from *t* + 1. With the cost matrix, Waddington-OT [18] is then used as the algorithm to estimate a transport matrix 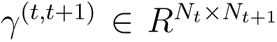. Each row in 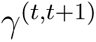 sums to 1, representing the transition probabilities of a cell in time step *t* to all the other cells in time step *t* + 1. Given *T* time steps, we need to maintain a total of *T −* 1 transport matrices throughout the autoencoder training process. For a given mini-batch *B* in each iteration, a submatrix version of 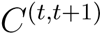 is computed using the rows and columns specified in *B* and is denoted by 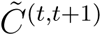. Similarly, a mini-batch version 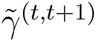 of 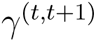 is calculated by taking the appropriate submatrix and rescaling the rows to unit sum. The batch-wise feature alignment loss (for each modality *s*) is defined as

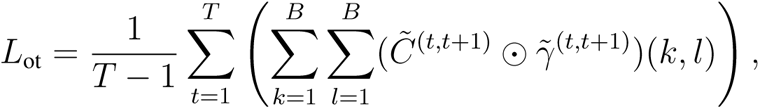

where 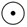 is the Hadamard product. The final *L*_ot_ is the sum over modalities *s*.
3. *Modality alignment loss*. For each mini-batch, the modality alignment loss is simply defined as the L2 distance between feature vectors from the same cell in the embedding space, which is to be minimized:

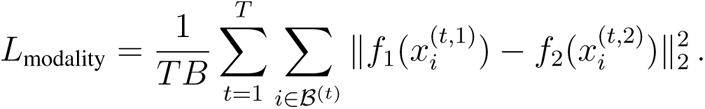

The total loss in each iteration is 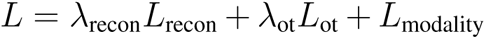 where the *λ*’s are tuning parameters controlling the relative weighting of the losses. For every *K* epochs, the transport matrices (for each modality *s*) 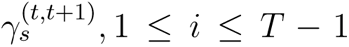 are updated by computing OT on the current embedding features.

### Training details

scTIE took a collection of peak matrices from scATAC-seq data and raw couns matrices from scRNA-seq data from multiple time points as input. For ATAC, the peak matrices were transformed to binary matrices, where one represents any non-zero original values. For RNA, the raw count matrices were sized-factor normalized and then log-transformed. For the overall multimodal training, we first pre-trained the RNA autoencoder *f*_1_*, g*_1_ for 500 epochs (excluding *L*_modality_). Then, we fixed the weights of the pretrained RNA model to train the ATAC model for 300 epochs with the overall loss *L*. Finally, the two models were jointly trained for 200 epochs using the full algorithm as detailed in Algorithm 1. The final joint embeddings were calculated by taking the averages of 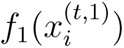 and 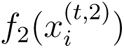 for each cell *i* from time *t*, followed by computing the final 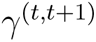 from the joint embeddings. Throughout training, we used Adam as the optimizer with learning rate set to 0.1, batch size *B* = 256, tuning parameters *λ*_recon_ = 1, *λ*_ot_ = 0.1, and OT was updated every 10 epochs.

**Algorithm 1.**
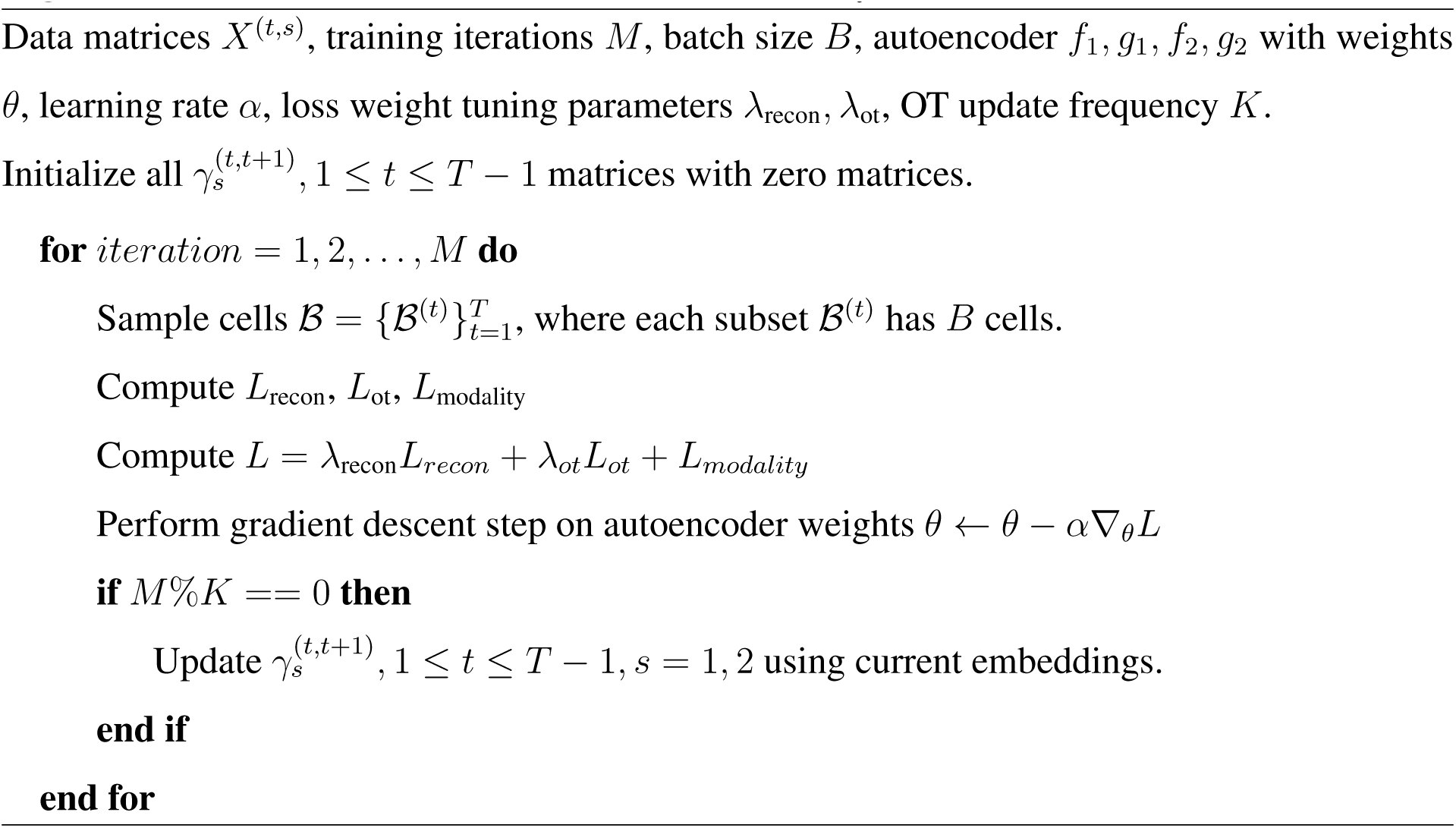
Multimodal OT Autoencoder (two-modality case)

### Cell type annotation of mESC data

#### Cell clustering of scTIE

To identify the clusters on the common embedding of scTIE, we first constructed a shared nearest neighbor graph using `buildSNNGraph` in R package `scran` [55] (v 1.23.0), with the number of nearest neighbor set as 15 with weighted scheme set as jaccard. Next we performed Leiden community detection [56] on the shared nearest graph with resolution 1.8 and number of iterations 50, implemented in R package `leidenAlg` (v 1.0.3), resulting in 17 clusters in total.

#### Motif enrichment

We used `Signac` [57] to calculate the over-represented motif of each cluster based on the differential accessible peaks. The motif position frequency matrices are obtained from cisBP [58]. We used `limma-trend` [59] to perform differential accessibility analysis between the cells in one cluster and the remaining cells, where the top 500 peaks of each cluster with log fold change greater than 0.1 and adjusted p-value less than 0.001 are selected. We then performed the motif enrichment analysis using `FindMotifs` to find motifs over-represented in the selected set of peaks.

### Benchmarking and evaluation metrics

#### Settings used in other methods

We benchmarked the performance of scTIE against four other methods designed for single-cell paired multimodal data integration: Seurat, scAI, MultiVI and MOFA. We compared scTIE’s performance in terms of visualisation of the latent space, alignment of the days and clustering in the latent space against these methods.

- **Seurat**. R package Seurat v4.1.0 [15] was used. We ran Seurat (WNN) using `FindMultiModalNeighbors`, with the reduction list input as the first 50 components of LSI reduced dimension of scATAC-seq (with the first dimension excluded) and 50 top PCs of scRNA-seq, with other parameters set as default.
- **scAI**. R package scAI v1.0.0 [16] was used. We ran scAI using `run scAI` by setting the rank of the inferred factor set as 64 and `nrun = 5`, with other parameters set as default.
- **MultiVI.** Python package scvi v0.15.0 [14] was used. We ran MultiVI using `MULTIVI` by setting the `fully paried = True`, `n hidden = 256` and `n latent = 64`, with other parameters set as default. The model was then trained with `max epochs = 200`.
- **MOFA**. R package MOFA2 v1.7.0 [13] was used. We ran MOFA using `run mofa` by setting the number of factors as 64, with other parameters set as default.

### Benchmarking of mESC data

#### Modality alignment

We used two metrics to measure scTIE’s performance in the alignment of the two modalities, namely FOSCTTM and paired data proportion.

- *FOSCTTM*. FOSCTTM refers to Fraction of Samples Closer than True Match, which is first introduced in MMD-MA [60] to quantify the alignment of multi-omics data. To evaluate the modal alignment of scTIE using FOSCTTM, we first calculated the Euclidean distance between the ATAC embedding and RNA embedding. Then for each modality we calculated one FOSCTTM score, which summarizes the proportion of cells that are closer to the ground truth matched cells based on the distance matrix. Finally we summarized the FOSCTTM scores from the two modalities into one score by taking the average.
- *Paired data proportion*. Paired data proportion (used in Cobolt [7]) calculated the proportion of cells whose ground truth matched cells are included within a certain number of neighbors, based on the Euclidean distance between the ATAC embedding and RNA embedding. We varied the number of neighbors from 1 to the total number of cells in the data.

#### Day alignment

We quantified the alignment of data sampled on different days using neighborhood purity using neighborPurity in R package bluster (v1.5.1), which calculated the proportion of cells from the same day among a certain number of neighbors, based on the UMAP coordinates generated from the common latent embeddings.

#### Comparison with single-modality clustering

We benchmarked clustering results from scTIE against other paired data integration methods by evaluating how similar the results are compared to clustering dimension-reduced scRNA-seq (PCA space) or scATAC-seq (LSI space) alone. On the latent space of each method or the dimension-reduced space from scRNA-seq or scATAC-seq, we performed Leiden clustering on the shared nearest neighbor graphs constructed, with the same parameter settings as mentioned in Section *Cell clustering*. Note that for Seurat, we performed Leiden clustering directly on the weighted nearest neighbor graph it outputs. We used two metrics to quantify the results, Adjusted Rand Index and silhouette coefficient.

- *Adjusted Rand Index (ARI)*. We computed the ARI scores of clustering results from each data integration method and clustering results from scRNA-seq or scATAC-seq alone.
- *Silhoutte coefficient*. For each clustering result, we computed the silhouette coefficient based on the Euclidean distance calculated from the UMAP coordinates generated from the dimension-reduced scRNA-seq or scATAC-seq.

For both metrics, higher values indicate a method better captures the clustering information in a single modality.

#### Benchmarking of synthetic data

We benchmarked the data integration performance of scTIE with the other paired data integration methods in terms of three evaluation metrics: (1) ARI scores of the cell type annotation provided by the original study and the Leiden clustering results from each method; (2) neighborhood purity of days; and (3) neighborhood purity of batch for scenarios with synthetic batch effects.

### Enrichment analysis for embedding dimensions

Upon completion of training, scTIE has projected the high dimensional feature vectors (genes and peaks) into a 64 dimensional embedding space. Treating each dimension as a representation unit, for each cell type, we backpropagate the gradient of each unit with respect to gene and peak input to select features with the largest impact. More specifically, for each cell in cell type *G*, we pass its gene expression vector through the autoencoder to obtain its embedding vector *y* and compute 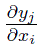 for each dimension *j* and gene input node *i*. The gradients are averaged over all cells in *G* to obtain the mean gradient for each gene. We then take the variability of gene expression into account by multiplying each mean gradient by its corresponding gene standard deviation, so that the final gradients are equivalent to gradients after the first batchnorm layer. Finally, we rank the genes by their gradient values and calculate the enrichment scores of the top 200 genes from the DE analysis of cell type *G*, where the DE analysis is performed using `limma-trend` [59] between the cells in one cluster and the remaining cells. Similar steps are performed for the peaks and the top 500 peaks are selected for enrichment score calculation.

We used *fgsea* function in the R package *fgsea* [61] to perform the gene set enrichment analysis (GSEA) on the pathways related to mouse embryonic stem cells (as listed in **Fig. 4B**). Significant pathways are defined with adjusted p-value less than 0.05.

### GRN inference

#### Selecting features with high predictive power

By building a prediction framework on the obtained transition probabilities, scTIE selects genes and peaks jointly with high predictive power for developmental outcomes. In the mESC data, we consider how a group of cells from earlier days, denoted as *G*_0_, develops into two other groups *G*_1_ and *G*_2_ on later days.

The transition probabilities are obtained from 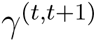 (*t* = 1, 2 in our data) so that each cell *i* in *G*_0_ is associated with a probability vector (*p_i_*_1_*, p_i_*_2_) indicating its probabilities of becoming *G*_1_ and *G*_2_ (See Section *Cell transition probability calculation*). We finetune a one-layer classifier on the pretrained features in the embedding space of cells in *G*_0_ to predict their transition probabilities. A simple linear classifier is sufficient to partition the cell feature space into *G*_1_ and *G*_2_ when the pretrained features are representative enough. Concretely, let *q* be the linear classifier and *B* be a mini-batch of cells from *G*_0_ of size *B*, we employ a batch-wise KL divergence loss defined

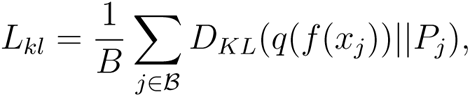

where *f* is the trained encoder, *P_j_*= (*p_j_*_1_*, p_j_*_2_). This loss enforces the classifier *q* to output transition probability distributions close to those in *P_j_*’s. We also include the modality alignment loss *L*_modality_, with weight default set as 0.1. The classifier is trained with Adam setting learning rate to 0.001, training epochs to 200, batch size to 256 and L1 regularization.

After training, gradients from the two classification nodes are backpropagated to each gene (or peak) input the same way as in computing embedding gradients. The gene gradients are then scaled by multiplying with the gene-wise standard deviations. A positive gradient for gene (or peak) *j* with respect to the node for *G*_1_ means increasing the input feature value tend to increase the cells’ probabilities of becoming *G*_1_, while a negative value indicates more contribution to *G*_2_. The final feature ranking is based on the average gradients by repeating this procedure 20 times with different seeds.

### Selection of *G*_0_*, G*_1_*, G*_2_

As a case study in this paper, we focus on the transition of cells from anterior primitive streak on Day 2 and Day 4 into endoderm, mesoderm, as well as remaining as anterior primitive streak on Day 4 and Day 6.

First, we considered the cells that are annotated as anterior primitive streak (Cluster 6) on Day 2 and Day 4 as *G*_0_. *G*_1_ and *G*_2_ are then selected from the cells on Day 4 and Day 6 that are more likely to be the descendants of *G*_0_, as quantified by the descendant scores. The descendant scores are defined similarly as in WOT [18]. Recall 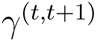 is the *N_t_*by *N_t_*_+1_ transition probability matrix between time points *t* and *t* + 1, let 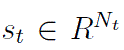 be the vector of descendant scores for all cells at time point *t*, then we can calculate

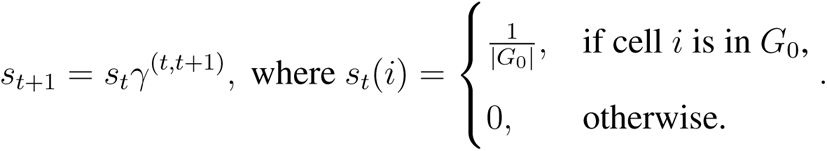

This formula can then be pushed forward again to calculate the descendant scores for the next time point *t* + 2, and so on. For all cells in *G*_0_ at time point *t* (here *t* = 1 or 2), we calculated the descendant scores *s_t_*_+_*_k_* of all cells at the later time point *t* + *k,* for *k* = 1*, …, T − t*. We then considered the cells with descendant scores greater than the median of all cells at a certain time point as the potential descendants, i.e., cells with *s_t_*_+_*_k_*(*i*) *>* median(*s_t_*_+_*_k_*). Among these descendant cells, we selected three pairs of *G*_1_ and *G*_2_ corresponding to the three cell fates we have analyzed: *G*_1_ that are annotated as (1) anterior primitive streak or (2) definitive endoderm or (3) mesoderm; for each selection of *G*_1_, *G*_2_ always represents the remaining descendant cells.

### Cell transition probability calculation

For each cell *i ∈ G*_0_ on Day *t*, and *G*_1_*, G*_2_ on Day *k ∈ K*, where *K* = *{k*: *t < k ≤ T }*, the transition probability vector 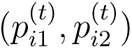 are calculated as the following,

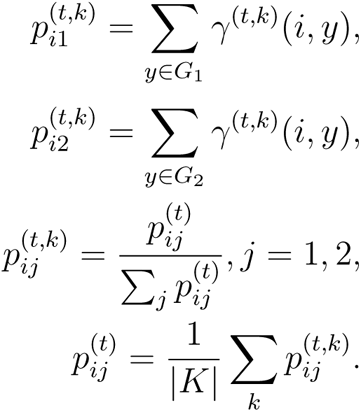

(*p_i_*_1_*, p_i_*_2_) is then the concatenated vector of 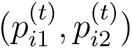.

### Evaluation of cell transition probability prediction

To evaluate the predictive power of the selected features to the transition probability, we performed support vector machine (SVM) with radial kernel to predict the transition probability using Day 2 and 4 anterior primitive streak gene expression of the top selected genes and peak matrix of the top selected peaks. The performance are quantified by root mean squared error (RMSE) from a 20 repeated 5 fold cross validation. We benchmarked the predictive power of the features selected by gradients with different regularization weights (0, 1, 10, 100), against the features selected by DE/DA analysis using limma-trend [59].

### Gene regulatory network construction

To construct the gene regulatory network for each cell fate (anterior primitive streak, definitive endoderm and mesoderm), we focus on the top 500 genes based on the gradient ranking. For each gene, we consider the open chromatin regions that are within 250kb upstream and downstream of its transcription start site (TSS) as well as ranked top 2000 according to the gradients as the distal candidate functional regions, which results in 396, 404 and 339 gene-peak pairs for the three cell fates respectively. We next filter the pairs based on the gene-peak correlation, calculated from the pseudo-cells. The pseudo-cells are constructed using the following strategies: We first randomly selected 100 cells from the anterior primitive streak cells on Day 2. For each cell, we looked for its 5 nearest neighbors based on the euclidean distances of the common embeddings. Then we calculate the Pearson correlation of the gene-peak pairs. This procedure is repeated 20 times and the gene-peak pairs with an absolute average correlation greater than 0.2 are retained (APS: 35, DE: 38 and MES: 17 pairs remained).

To link the peak region with the TF, we identified the enriched TF using *matchMotifs* function in R package *motifmatchr* of the peaks from the selected gene-peak pairs based on CIS-BP database [58]. We only consider if the TF are the top 500 genes. Finally, by linking the TF-region and peak-gene relationships, we construct the TF-gene regulatory networks that are associated cell fate probabilities.

## Declarations

### Availability of data and materials

All the raw and processed data produced in this study will be deposited in GEO database. scTIE was implemented using PyTorch (version 1.9.1) with code available at https://github.com/SydneyBioX/scTIE.

## Acknowledgements

We would like to thank Michael Blanco and Dhananjay Wagh from Stanford Genomics Service Center (SGSC) for their kind help on the preparation of 10x Genomics single cell multiome libraries. We also want to thank Xuhuai Ji form SGSC for providing sequencing services.

## Fundings

The Illumina HiSeq 4000 was purchased using a NIH S10 Shared Instrumentation Grant (S10OD018220). The Illumina NovaSeq 6000 was also purchased using a NIH S10 Shared Instrumentation Grant (1S10OD02521201). The authors gratefully acknowledge the following funding sources: Research Training Program Tuition Fee Offset and Stipend Scholarship and Chen Family Research Scholarship to Y.L.; AIR@innoHK programme of the Innovation and Technology Commission of Hong Kong to J.Y.H.Y. and Y.L.; the UT Austin Harrington Faculty Fellowship to Y.X.R.W; NIH grants R01 HG010359 and P50 HG007735 to W.H.W.

## Author contributions

T.W., W.H.W. and Y.X.R.W. conceived and designed this project; X.C. performed the mESC multiome experiment; Y.L., T.W., S.W., B.C., and J.X. performed data preprocessing, model development, and evaluation of results; J.Y.H.Y., W.H.W. and Y.X.R.W. supervised the execution; Y.L., B.C., J.X., J.Y.H.Y., W.H.W. and Y.X.R.W. wrote the manuscript. All authors read and approved the manuscript.

## Competing interests

The authors declare that they have no conflict of interest.

## Supplementary materials

**Supplementary Figure S1:**
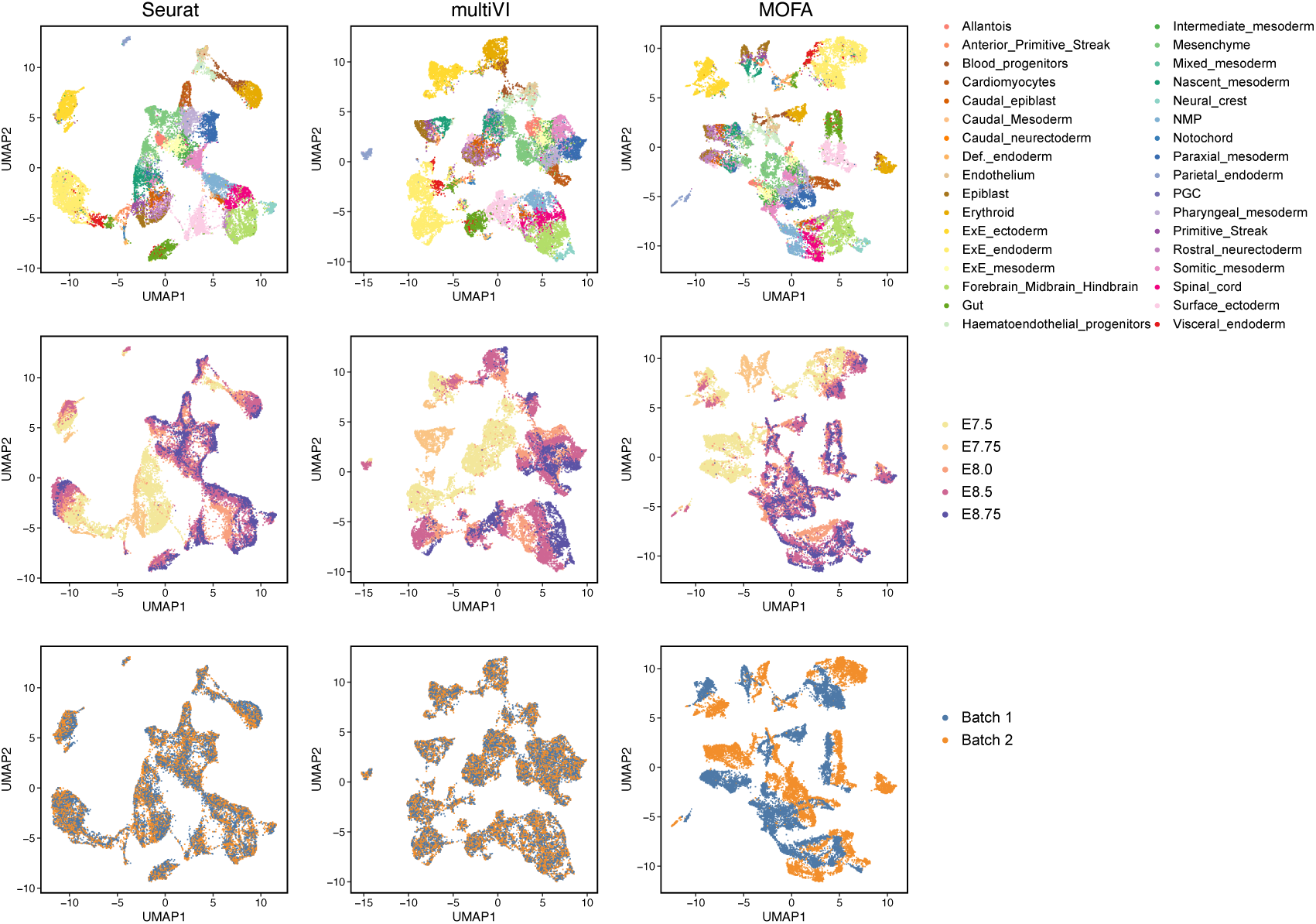
Joint visualization using UMAP of the synthetic dataset with batch effect in RNA and noise in ATAC for three data integration methods (Seurat, multiVI and MOFA), colored by cell type annotations (first row), sampling day (second row) and synthetic batch information (third row). Each dot represents a cell in the embedding space.

**Supplementary Figure S2:**
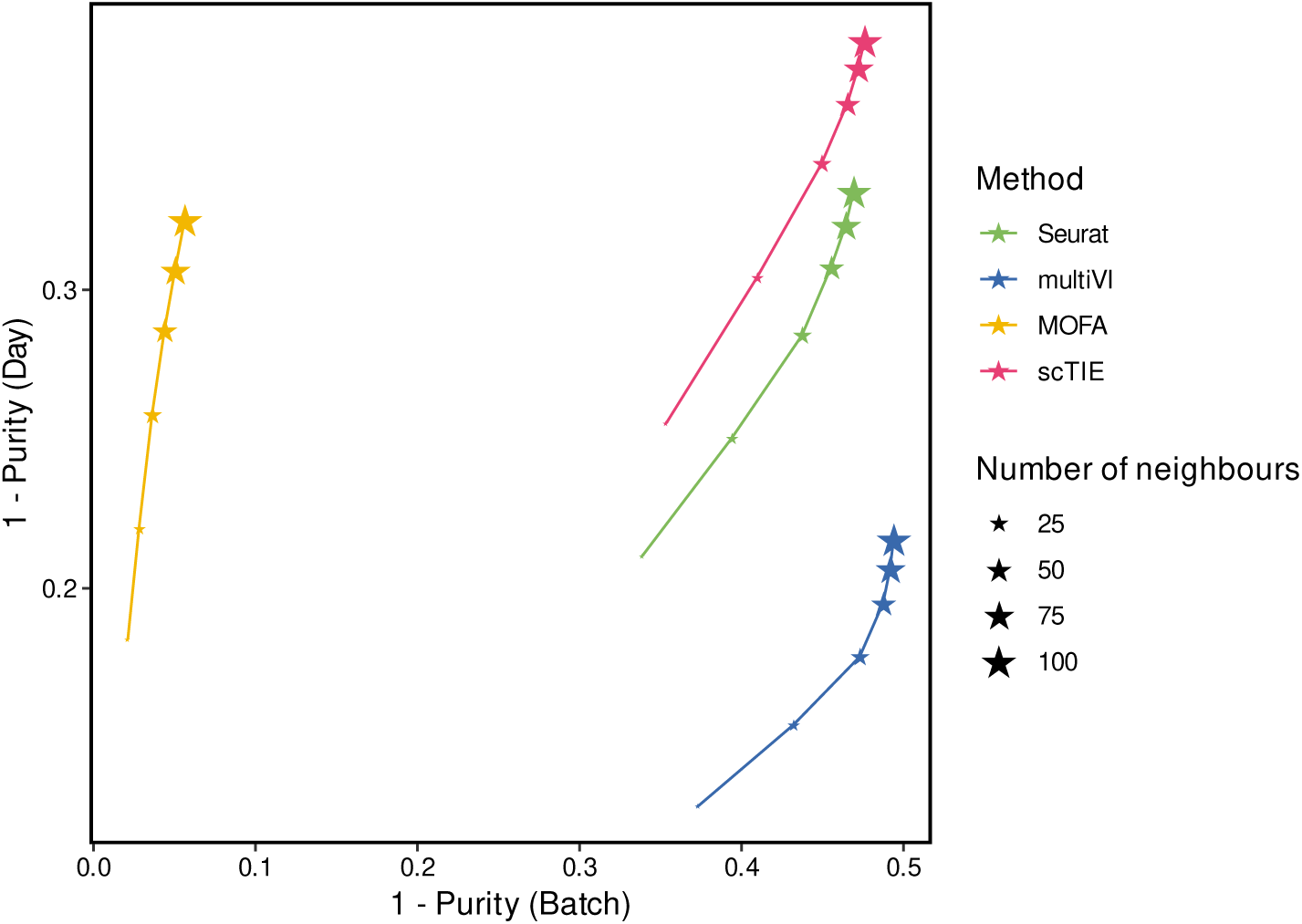
Scatter plot showing 1 – average purity scores of batch (x-axis) versus 1 – average purity scores of sampling day (y-axis) as the number of neighbors changes, where the size of stars represents the number of neighbors and color of the stars represents the method. Points in the top right corner have better day alignment and batch mixing.

**Supplementary Figure S3:**
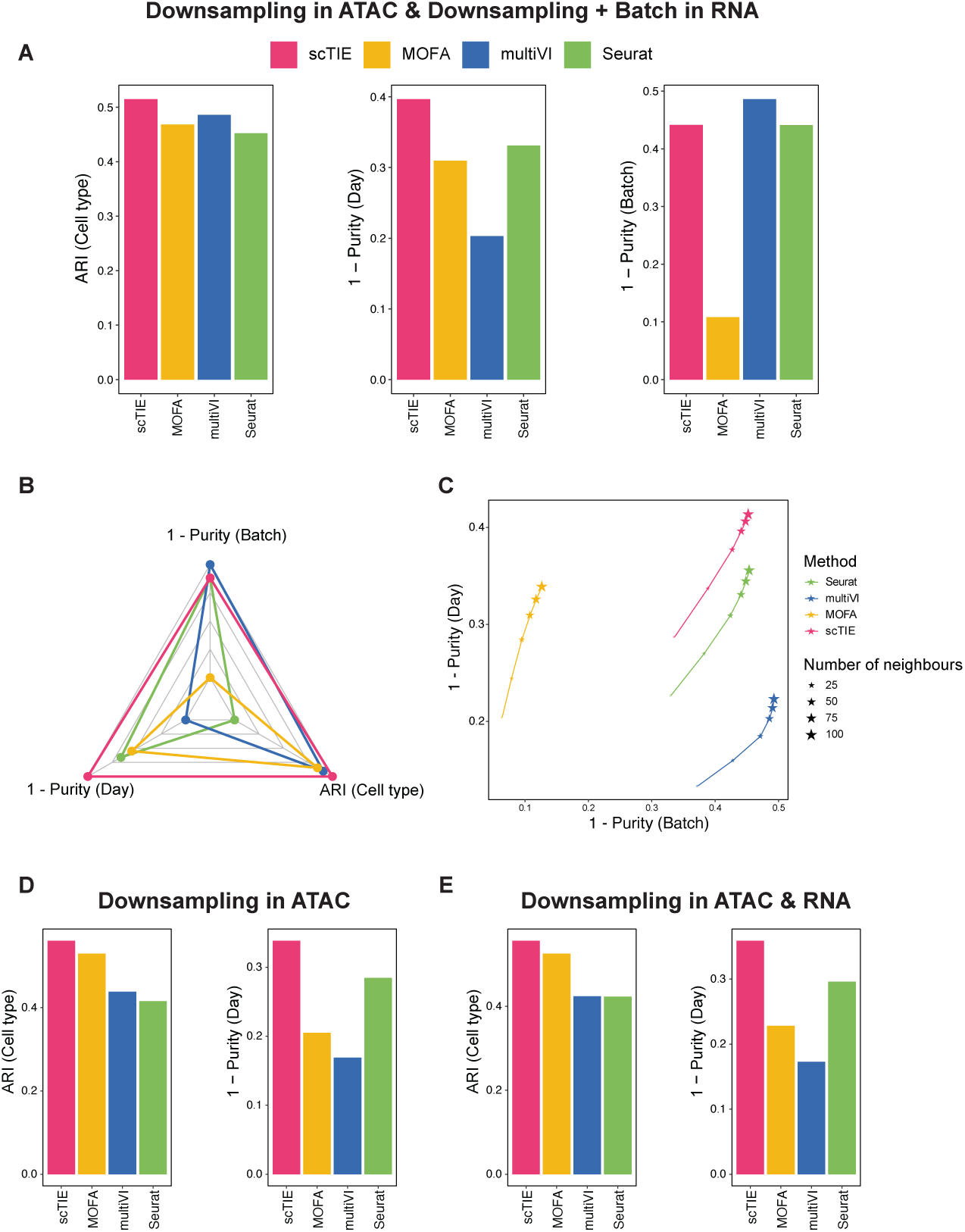
Evaluation results for variations of synthetic data settings: (A-C) Read downsampling in ATAC & Read downsampling + Batch effect in RNA: (A) Bar plots showing the evaluation metrics of different data integration methods, including ARI values for clustering with annotations (left); 1 – average purity scores of sampling day (middle) and 1 – average purity scores of the synthetic batch (right). (B) Radar plot summarizing the three evaluation metrics shown in (A), where each line represents the performance of one method, and each axis represents an evaluation metric, starting from the minimum value of all methods. (C) Scatter plot showing 1 – average purity scores of batch (x-axis) versus 1 – average purity scores of sampling day (y-axis) as the number of neighbors changes, where the size of stars represents the number of neighbors and color of the stars represents the method. (D-E) Bar plots showing the evaluation metrics of different data integration methods, including ARI values for clustering with annotations (left); 1 – average purity scores of sampling day (right) for (D) Read downsampling in ATAC and (E) Read downsampling in both ATAC and RNA.

**Supplementary Figure S4:**
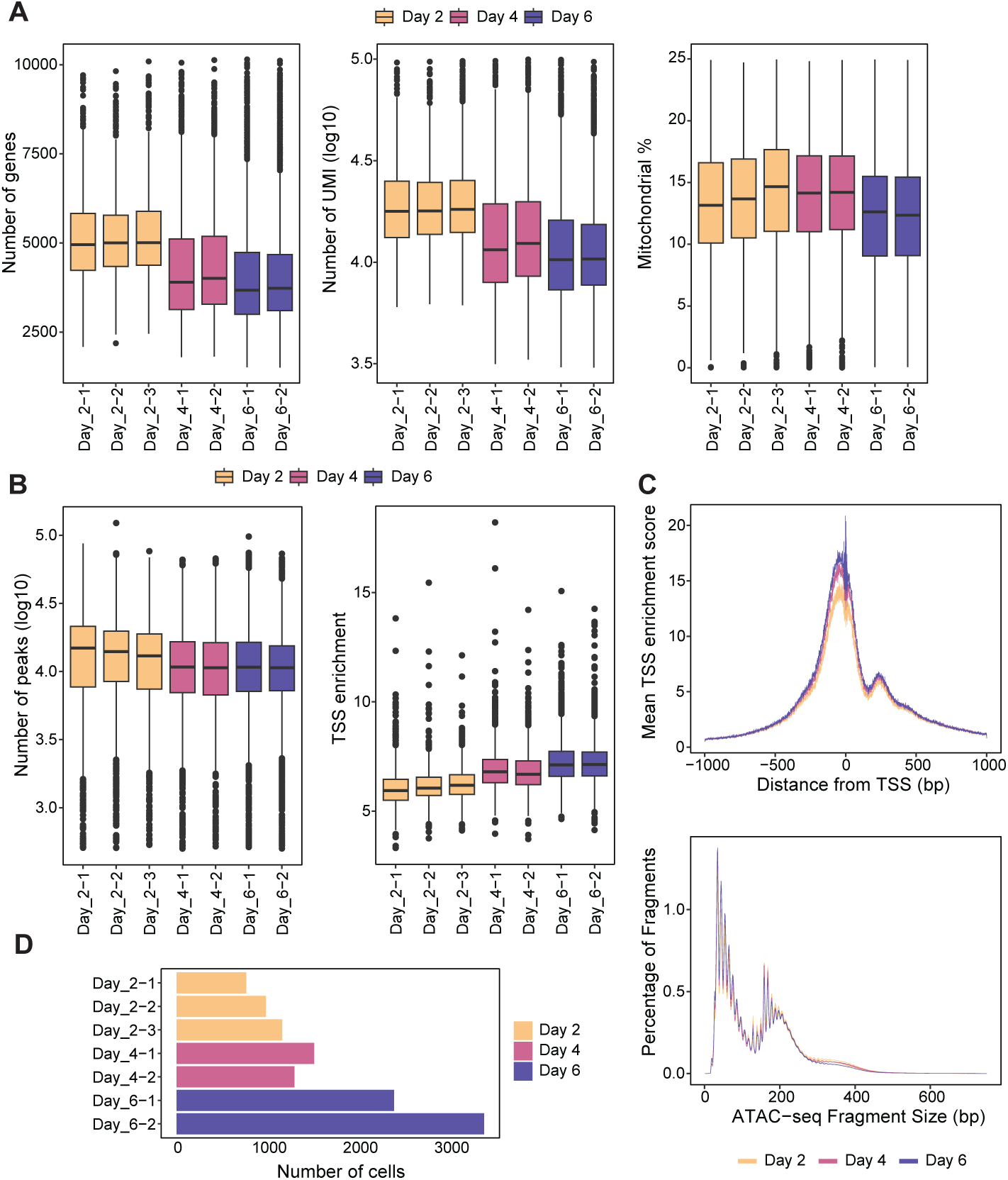
(A) Box plots showing the distribution of RNA quality metrics of each sample, color by the sampling day, including number of genes detected (left), number of total UMI (log10) (middle) and Mitochondrial (MT) gene fraction per cell (right). (B) Box plots showing the distribution of ATAC quality metrics of each sample, color by the sampling day, including number of peaks detected (log10) (left) and transcription start site (TSS) enrichment (middle). (C) Line plot showing the distribution of ATAC quality metrics of each sample, colored by sampling day, including normalized TSS enrichment score of each sample at each position relative to the TSS (first row) and fragment size distribution (second row). (D) Bar plots indicates the number of cells after quality control in each sample, colored by sampling day.

**Supplementary Figure S5:**
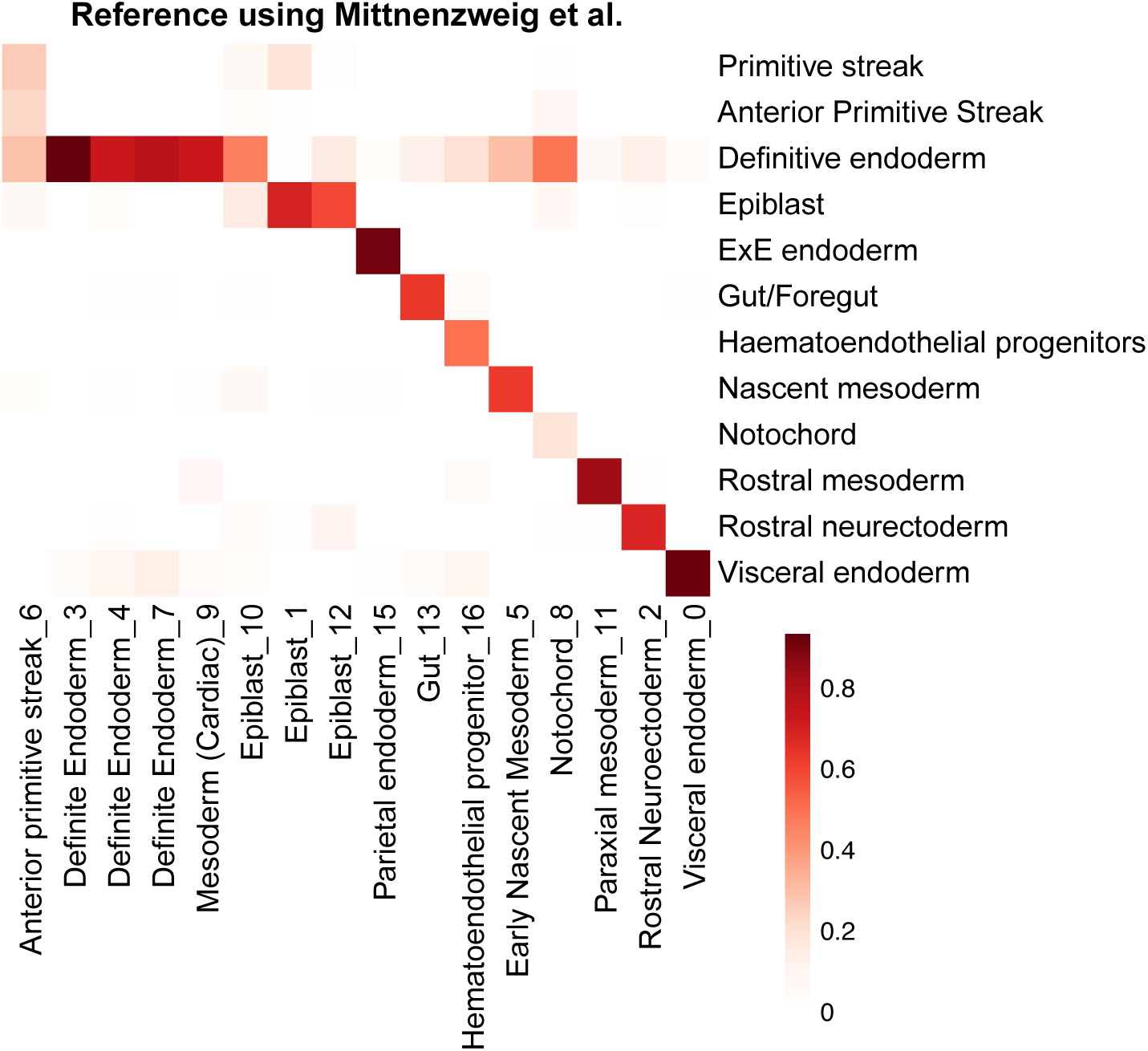
Heatmap comparing the clustering results and the transferred labels by scClassify [25] using Mittnenzweig data as reference [23]. Color indicates the proportion of cells classified as a certain cell type label in the reference for one cluster.

**Supplementary Figure S6:**
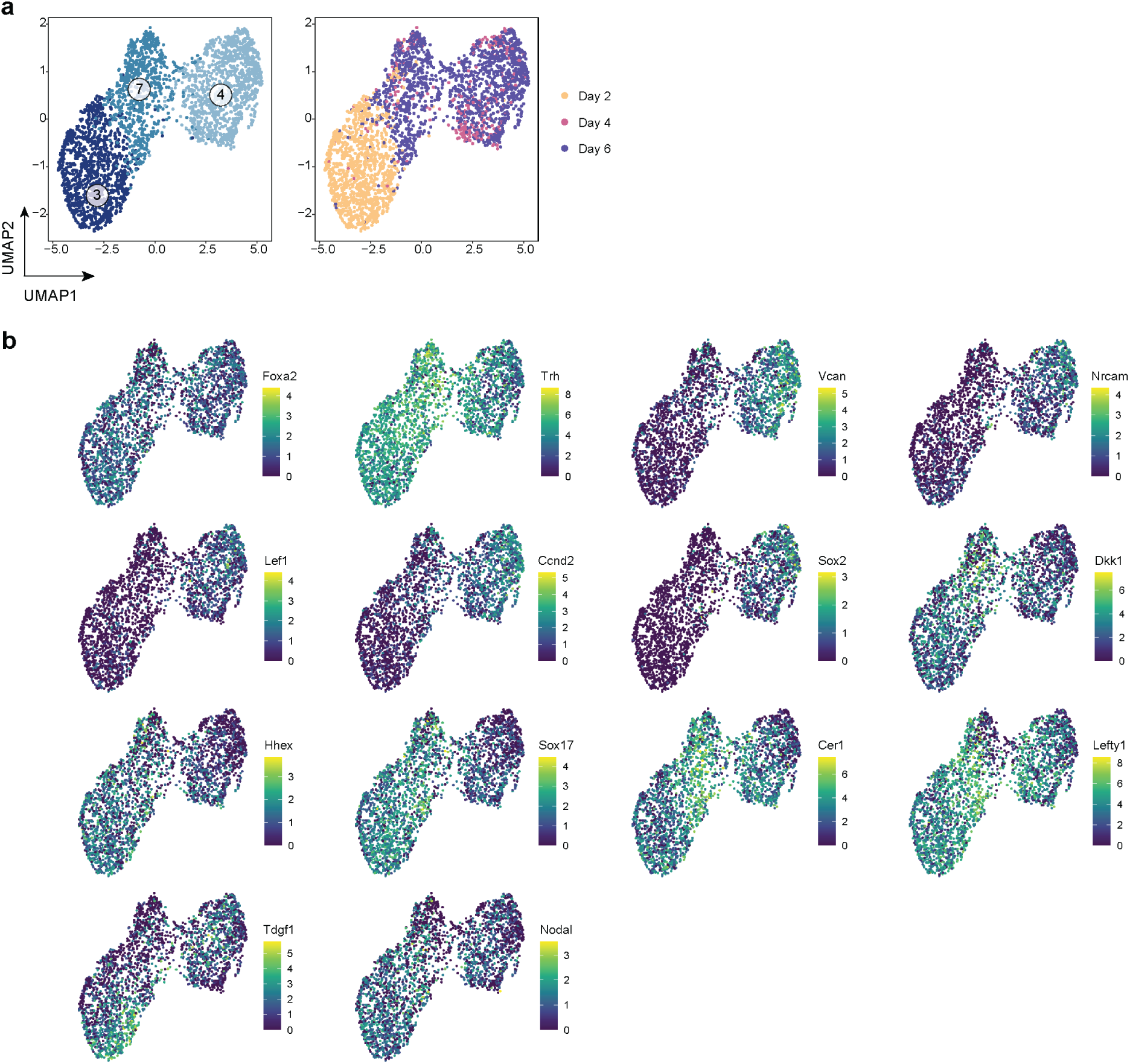
(A) UMAP of definitive endoderm, colored by clustering results (left) and sampling day (right). (B) UMAP visualisations of 14 selected markers of clusters.

**Supplementary Figure S7:**
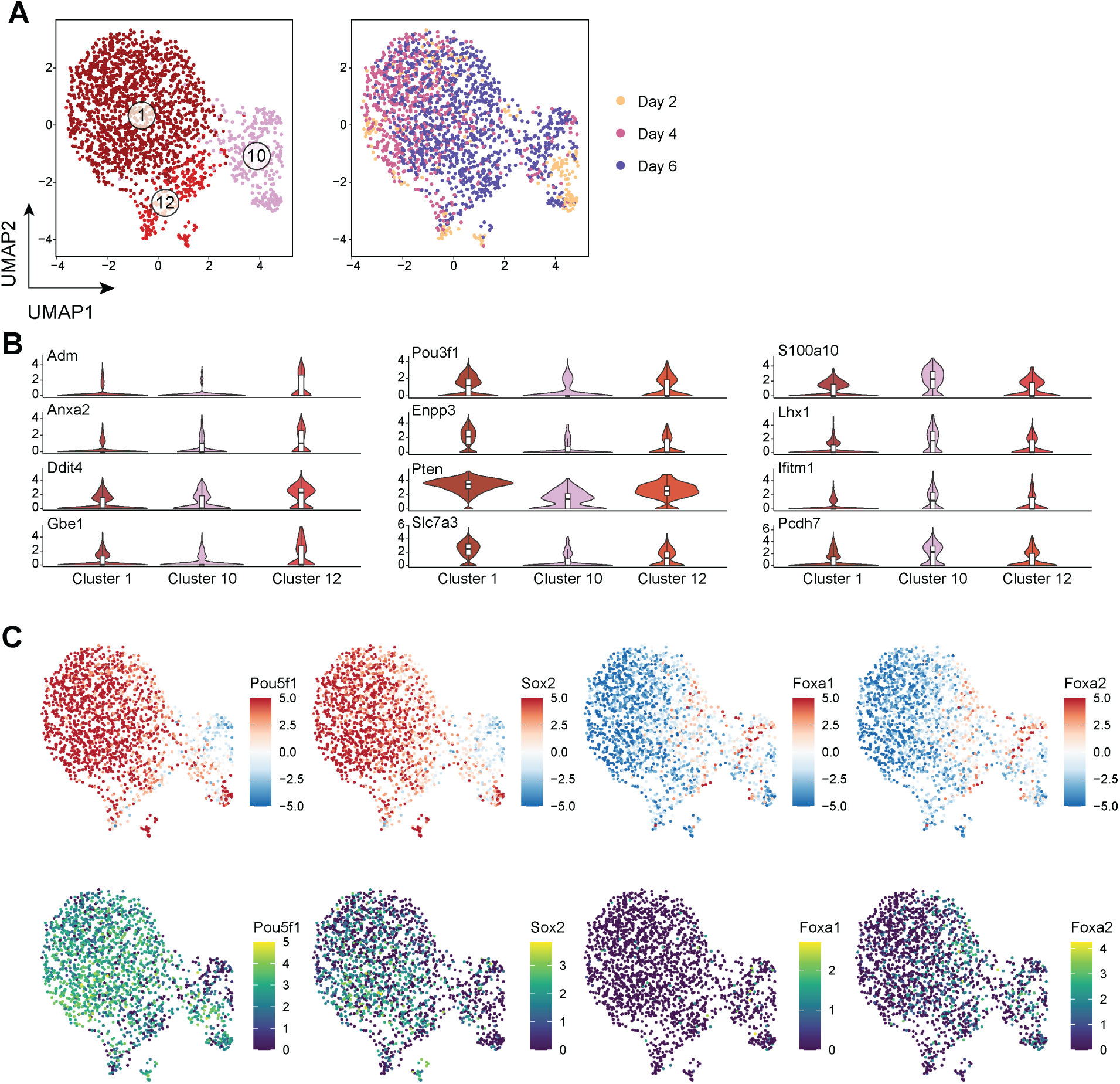
(A) UMAP of epiblast, colored by clustering results (left) and sampling day (right). (B) Violin plots showing the RNA expression of 12 selected markers. (C) UMAP visualisation of 4 selected TF, higlighted by the motif enrichment scores derived from ATAC using chromVar (top row) and RNA expression (bottom row).

**Supplementary Figure S8:**
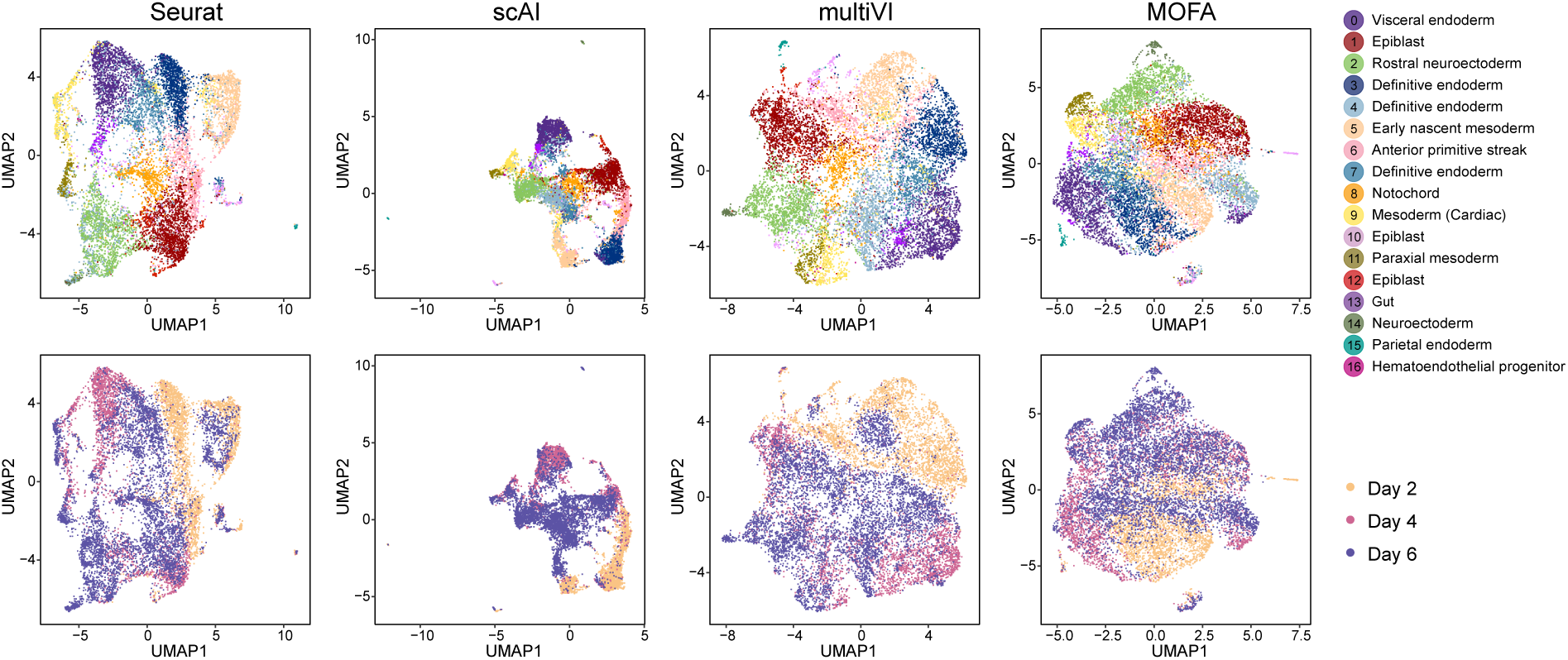
UMAP visualization of the dataset for Seurat, scAI, multiVI and MOFA, colored by annotated cell types (first row) and sampling days (second row).

**Supplementary Figure S9:**
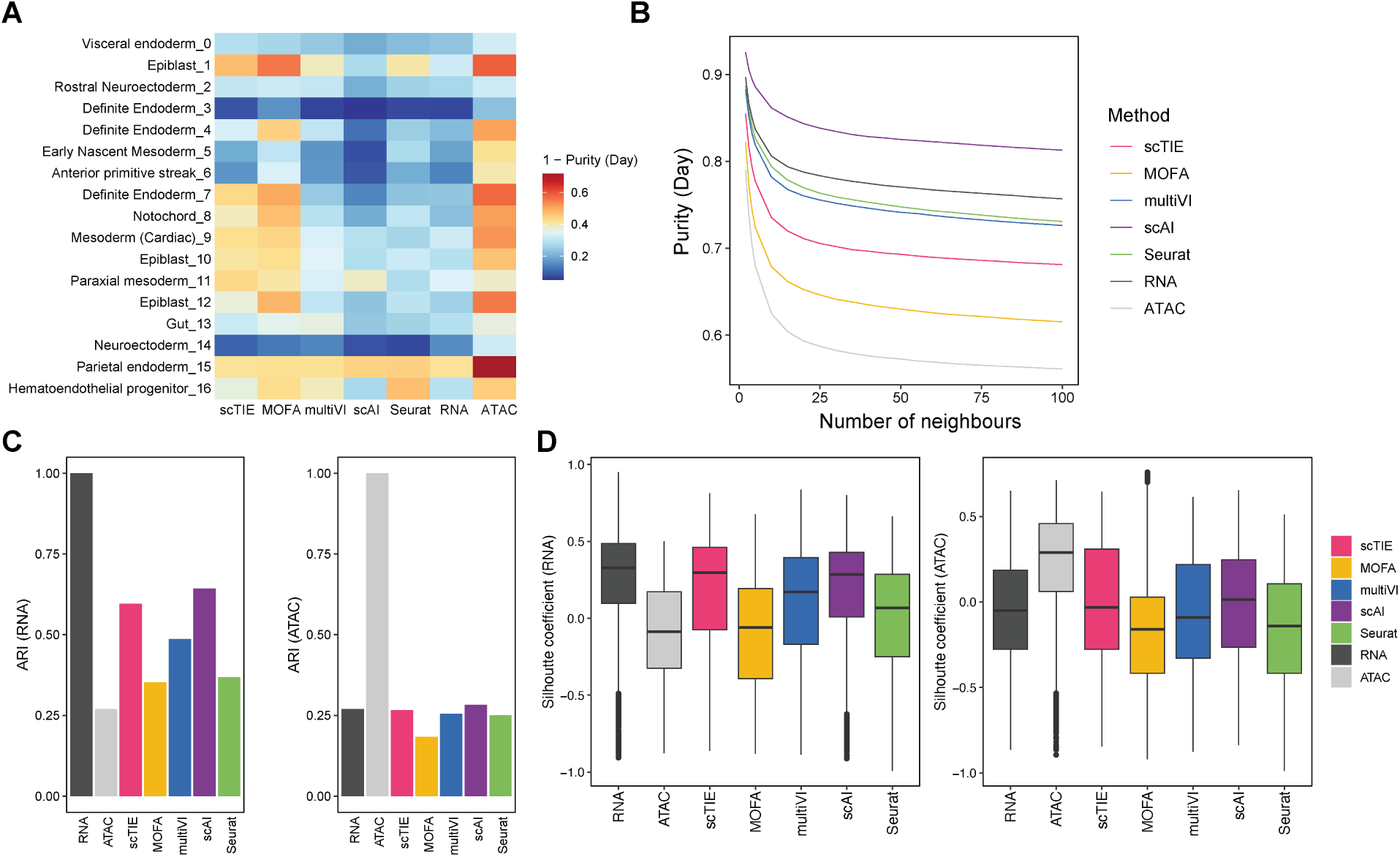
(A) 1 – Average purity scores of sampling day for each cell type (Number of neighbors = 50). Row indicates cell types and column indicates methods. Higher values indicate better mixing between days. (B) Average purity scores of sampling day based on different number of neighbors, colored by different methods. Lower values indicate better mixing between days. (C) The bar plots show the ARI values comparing the clustering from different data integration methods with clustering on RNA (left) and ATAC alone (right); higher values indicate better agreement. Note that here RNA and ATAC clustering results are the ground truth for the left panel and right panel respectively, therefore they have ARI equal to 1. (D) Box plots show the silhouette coefficient comparing the clustering from different data integration methods based on distance matrices computed from the RNA (left) and ATAC (right) UMAP coordinates. Higher values indicate better agreement. Note that for the left panel, RNA clustering result has the highest silhouette coefficients because clustering derived from RNA is used as the ground truth; similarly for the right panel.

**Supplementary Figure S10:**
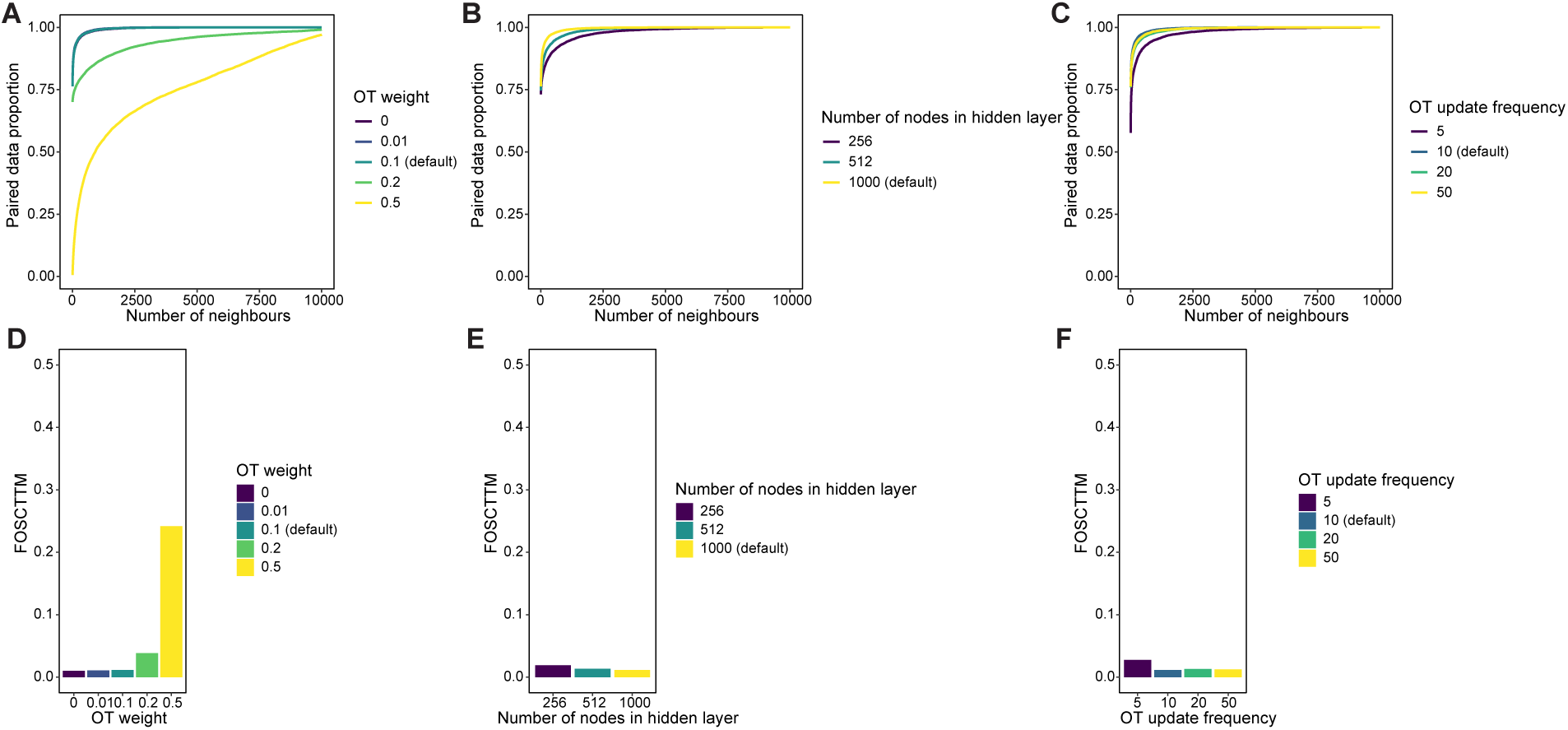
Robustness of scTIE with respect to the tuning parameters in modality alignment. (A-C) Proportion of ground truth pairs within certain number of nearest neighbors, with different (A) OT weight; (B) Number of nodes in hidden layer; (C) OT update frequency. (D-F) Barplots of FOSCTTM (fraction of samples closer than the true match) values, with different (D) OT weight; (E) Number of nodes in hidden layer; (F) OT update frequency.

**Supplementary Figure S11:**
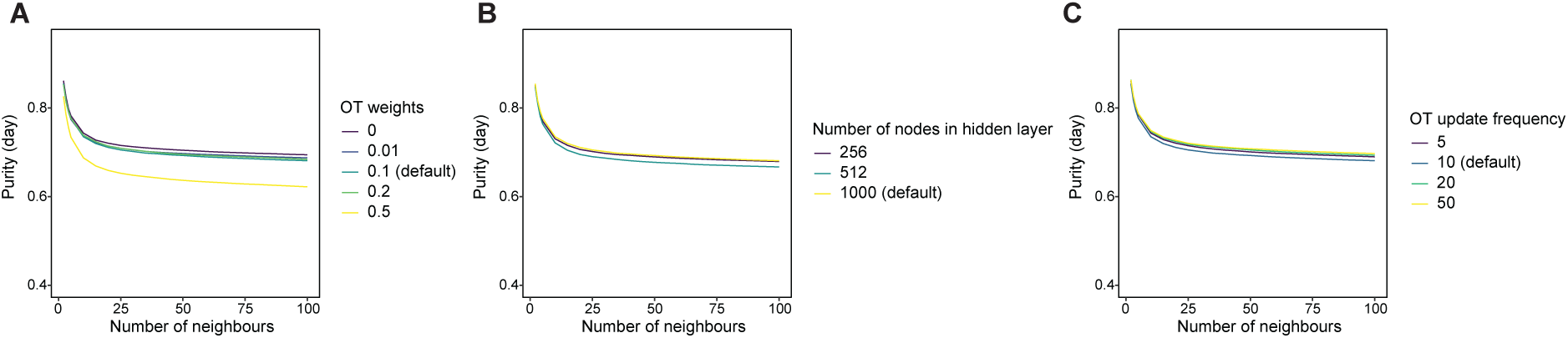
Robustness of scTIE with respect to the tuning parameters in time point alignment. Average purity scores of sampling days based on different number of neighbors, with different (A) OT weight; (B) number of nodes in the hidden layer; (C) OT updating frequency.

**Supplementary Figure S12:**
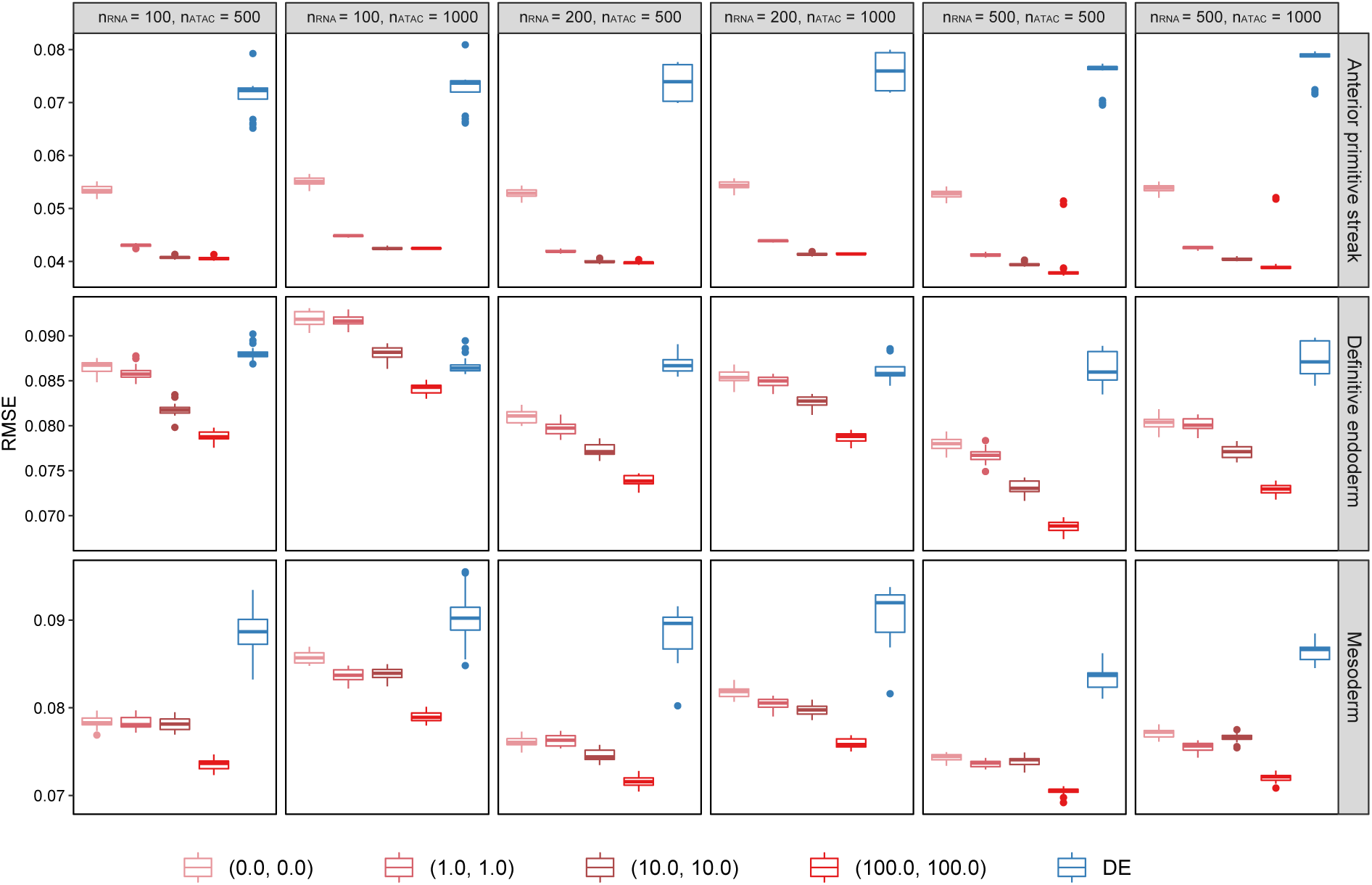
Evaluation of transition probability predictions for three different cell fates: anterior primitive streak, definitive endoderm and mesoderm, comparing (1) different number of genes/peaks; (2) different L1 regularization weights used in the prediction task.

**Supplementary Figure S13:**
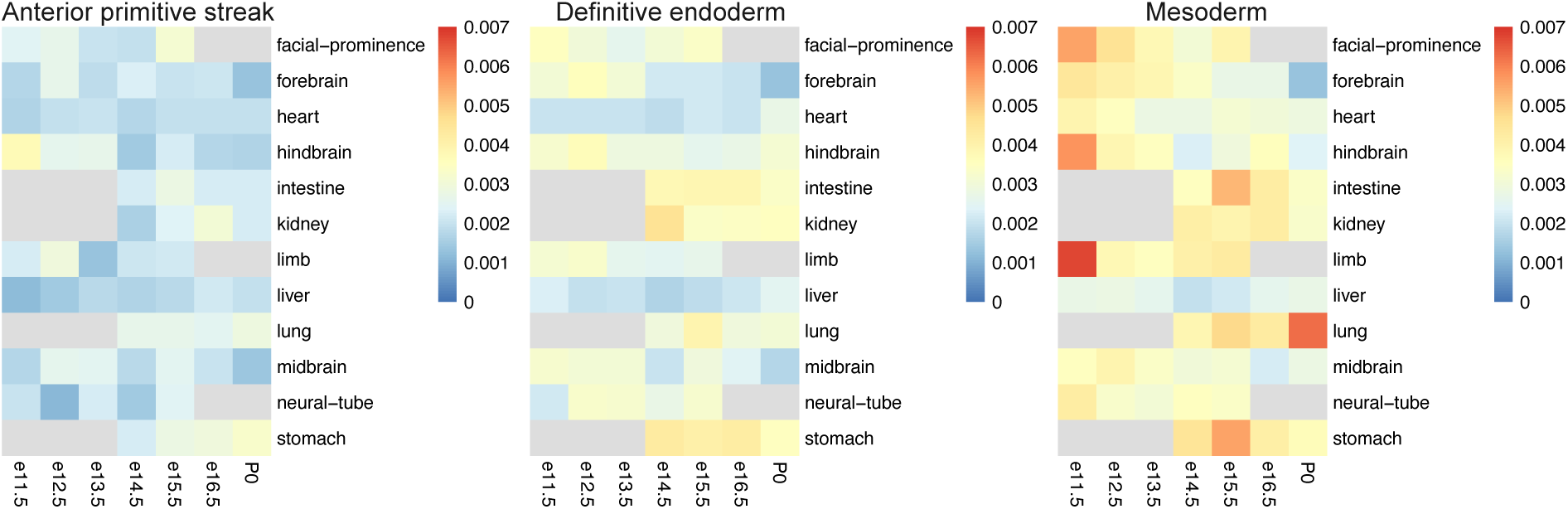
Similarity of top DA regions with enhancers of 12 tissues at seven developmental stages from known enhancer databases.

**Supplementary Figure S14:**
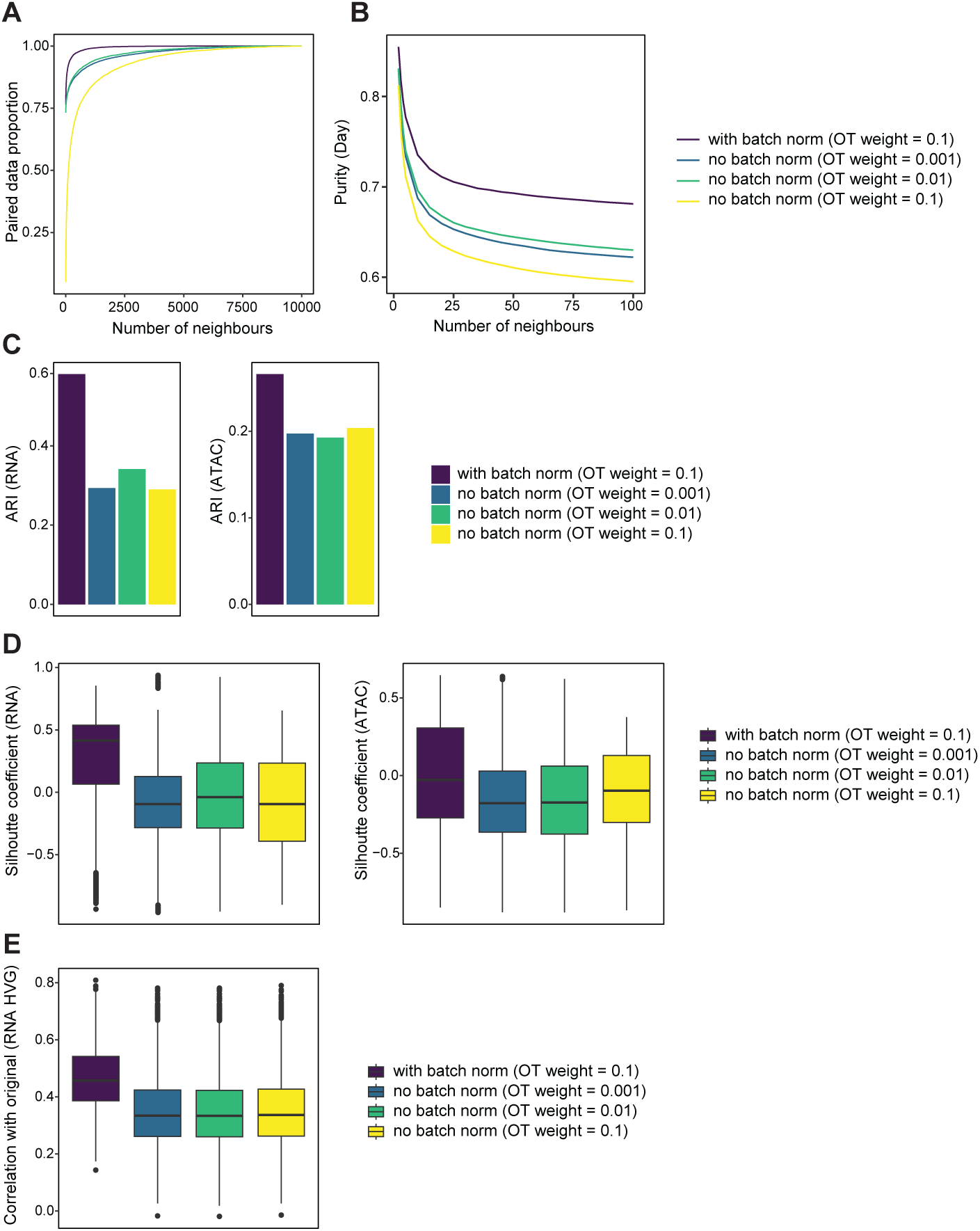
scTIE performance comparison with and without the coupled batch norm layers in RNA: (A) Proportion of ground truth pairs within a given number of nearest neighbors; (B) Average purity scores of sampling days based on different numbers of nearest neighbors; (C) The bar plots show the ARI values comparing the clustering from different settings of scTIE with the clustering on RNA (left) and ATAC alone (right); higher values indicate better agreement. (D) Box plots show the silhouette coefficients comparing the clustering from different settings of scTIE based on distance matrices computed from the RNA (left) and ATAC (right) UMAP coordinates. Higher values indicate better agreement. (E) Correlation of scTIE reconstructed RNA expression with the original RNA expression of highly variable genes (HVG).

**Table S1:**
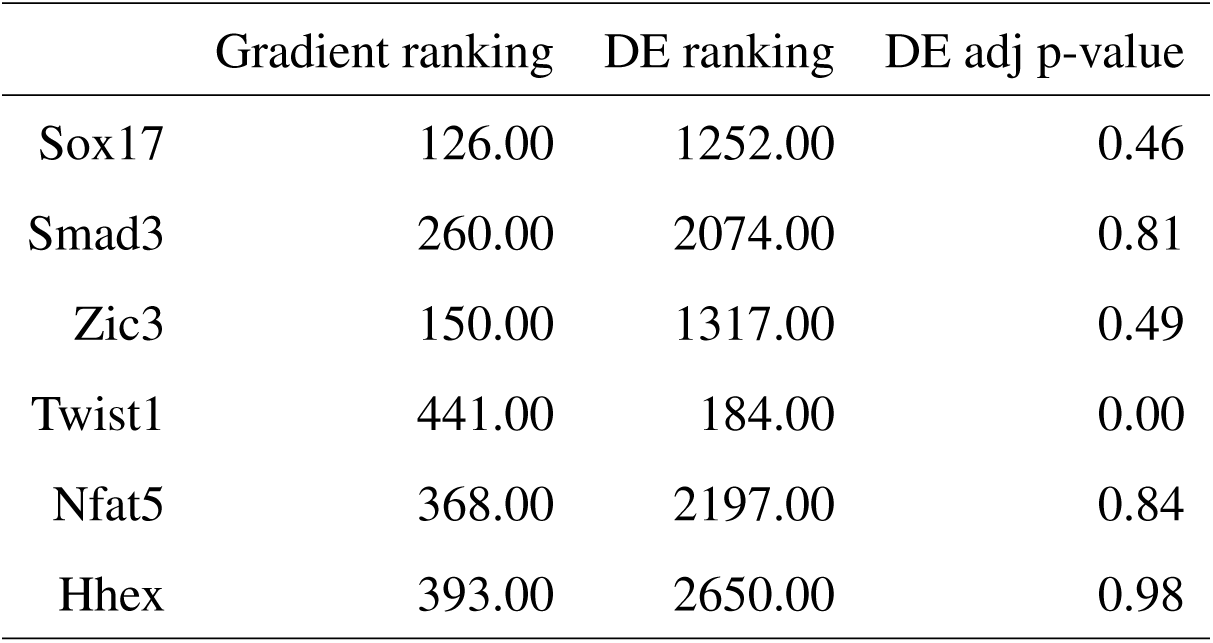
Comparison of the gradient rankings, DE rankings and adjusted p-values under DE for key TFs in mesoderm lineage.

## Notes

### Competing Interest Statement

The authors have declared no competing interest.

