## Supplementary figures and tables for "scTIE: data integration and inference of gene regulation using single-cell temporal multimodal data"

### Supplementary materials

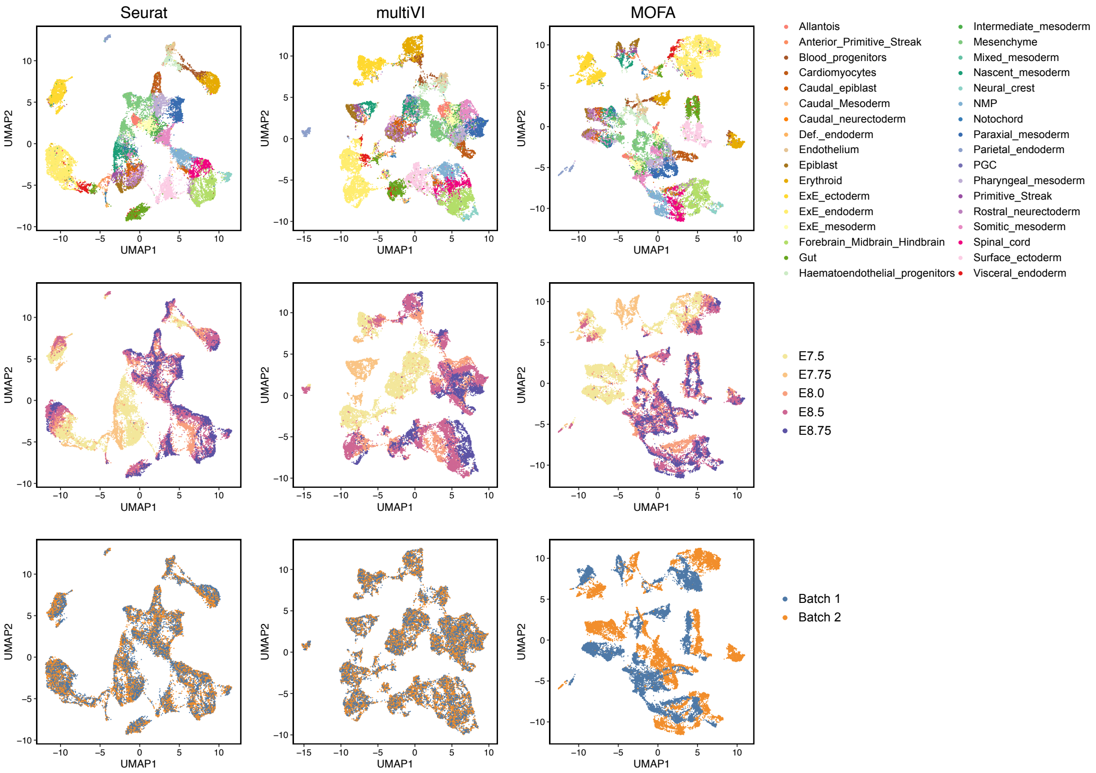


Supplementary Figure S1: Joint visualization using UMAP of the synthetic dataset with batch effect in RNA and noise in ATAC for three data integration methods (Seurat, multiVI and MOFA), colored by cell type annotations (first row), sampling day (second row) and synthetic batch information (third row). Each dot represents a cell in the embedding space.

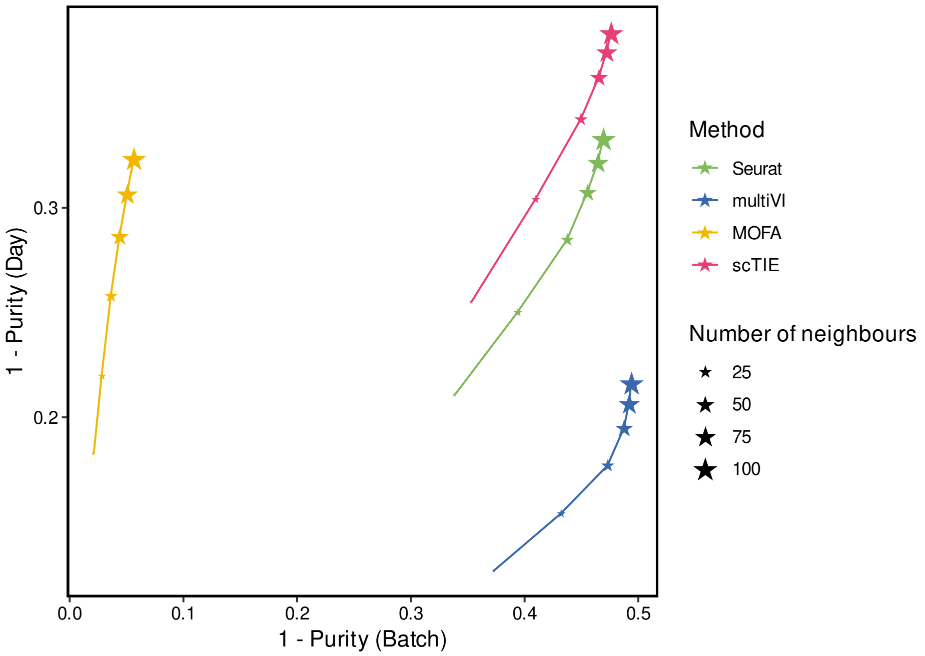


Supplementary Figure S2: Scatter plot showing 1 - average purity scores of batch (x-axis) versus 1 - average purity scores of sampling day (y-axis) as the number of neighbors changes, where the size of stars represents the number of neighbors and color of the stars represents the method. Points in the top right corner have better day alignment and batch mixing.

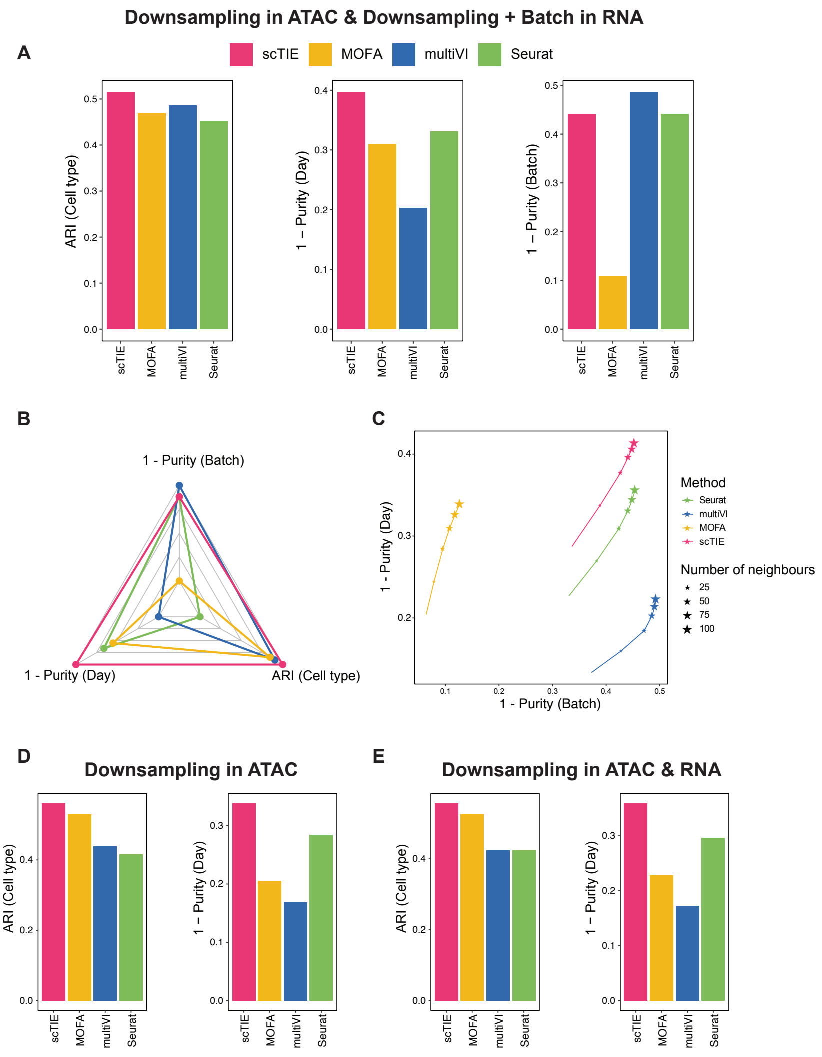


Supplementary Figure S3: Evaluation results for variations of synthetic data settings: (A-C) Read downsampling in ATAC & Read downsampling + Batch effect in RNA: (A) Bar plots showing the evaluation metrics of different data integration methods, including ARI values for clustering with annotations (left); 1 - average purity scores of sampling day (middle) and 1 - average purity scores of the synthetic batch (right). (B) Radar plot summarizing the three evaluation metrics shown in (A), where each line represents the performance of one method, and each axis represents an evaluation metric, starting from the minimum value of all methods. (C) Scatter plot showing 1 - average purity scores of batch (x-axis) versus 1 - average purity scores of sampling day (y-axis) as the number of neighbors changes, where the size of stars represents the number of neighbors and color of the stars represents the method. (D-E) Bar plots showing the evaluation metrics of different data integration methods, including ARI values for clustering with annotations (left); 1 - average purity scores of sampling day (right) for (D) Read downsampling in ATAC and (E) Read downsampling in both ATAC and RNA.

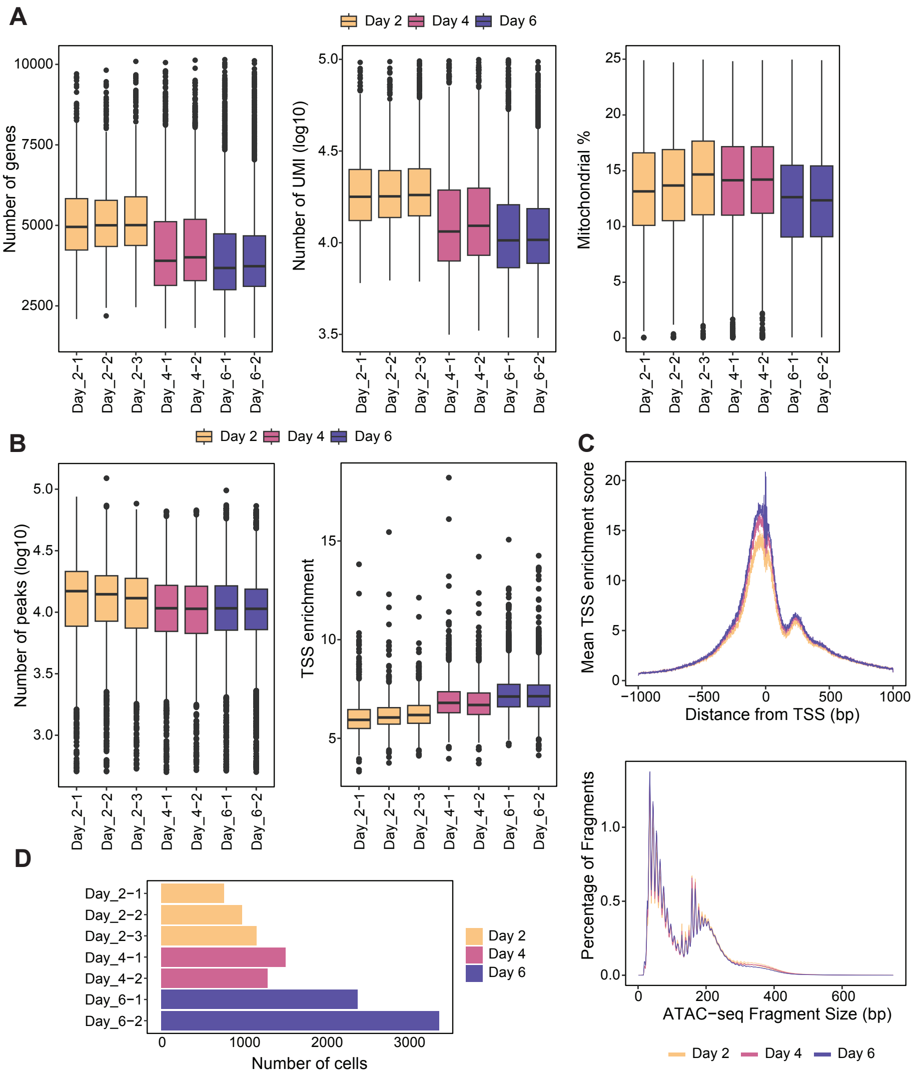


Supplementary Figure S4: (A) Box plots showing the distribution of RNA quality metrics of each sample, color by the sampling day, including number of genes detected (left), number of total UMI (log10) (middle) and Mitochondrial (MT) gene fraction per cell (right). (B) Box plots showing the distribution of ATAC quality metrics of each sample, color by the sampling day, including number of peaks detected (log10) (left) and transcription start site (TSS) enrichment (middle). (C) Line plot showing the distribution of ATAC quality metrics of each sample, colored by sampling day, including normalized TSS enrichment score of each sample at each position relative to the TSS (first row) and fragment size distribution (second row). (D) Bar plots indicates the number of cells after quality control in each sample, colored by sampling day.

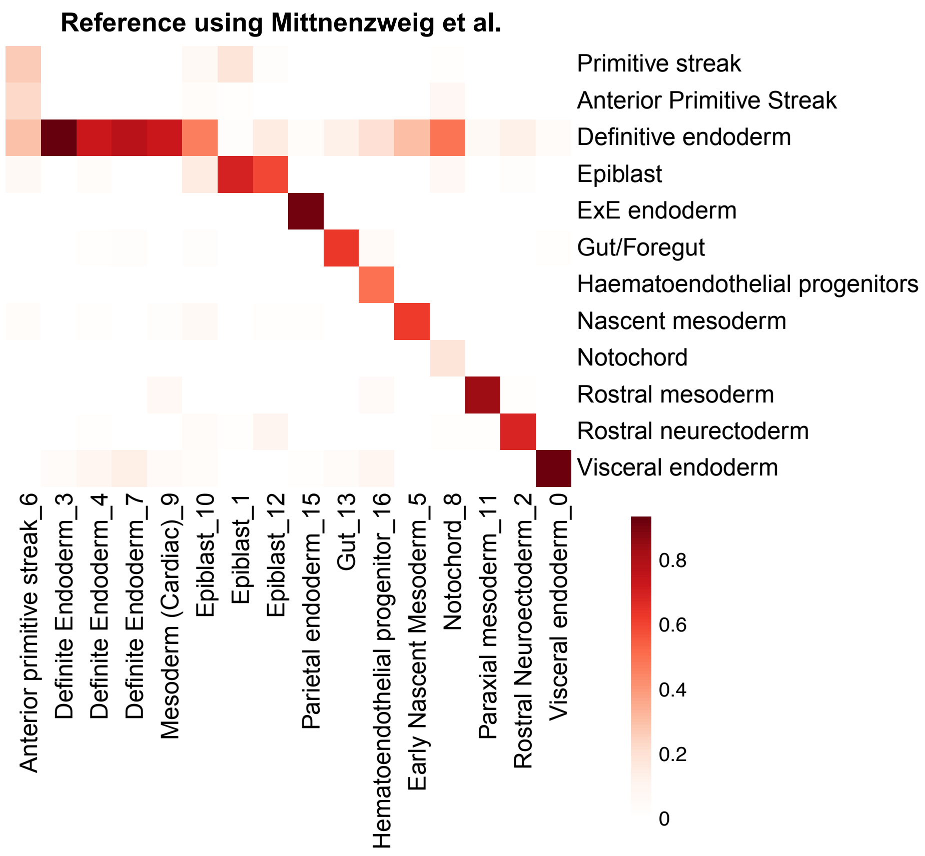


Supplementary Figure S5: Heatmap comparing the clustering results and the transferred labels by scClassify [25] using Mittnenzweig data as reference [23]. Color indicates the proportion of cells classified as a certain cell type label in the reference for one cluster.

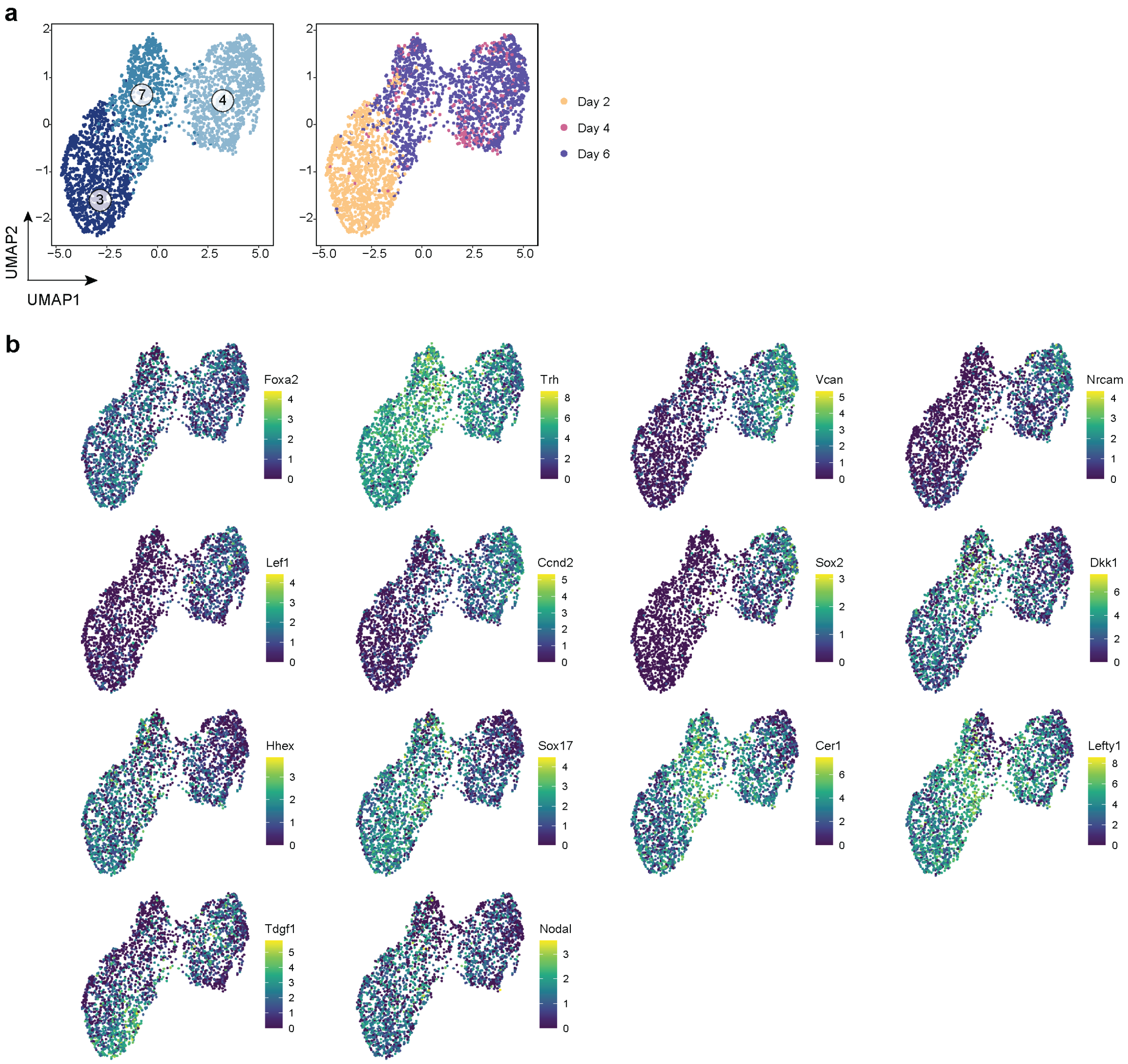


Supplementary Figure S6: (A) UMAP of definitive endoderm, colored by clustering results (left) and sampling day (right). (B) UMAP visualisations of 14 selected markers of clusters.

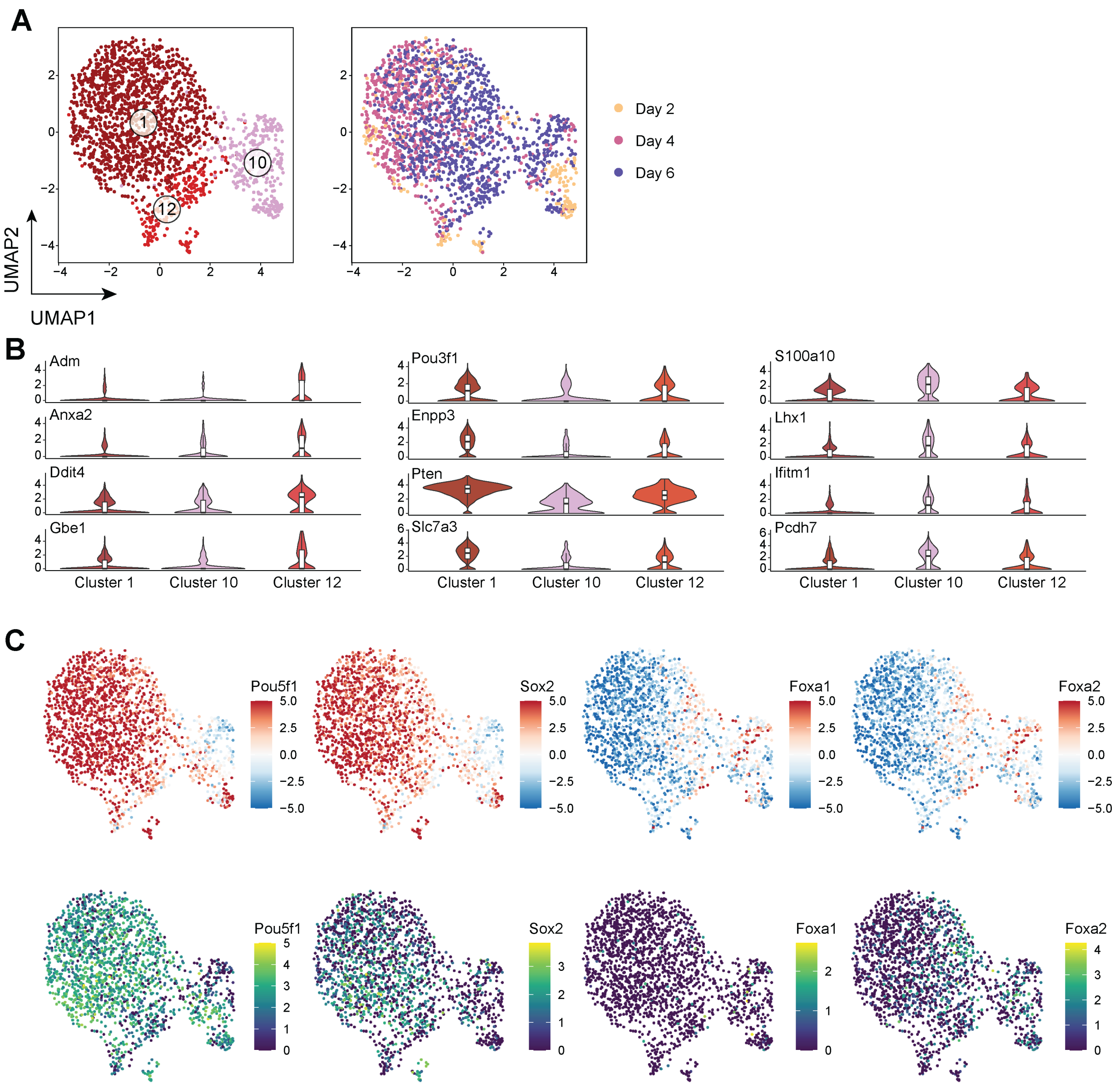


Supplementary Figure S7: (A) UMAP of epiblast, colored by clustering results (left) and sampling day (right). (B) Violin plots showing the RNA expression of 12 selected markers. (C) UMAP visualisation of 4 selected TF, higlighted by the motif enrichment scores derived from ATAC using chromVar (top row) and RNA expression (bottom row).

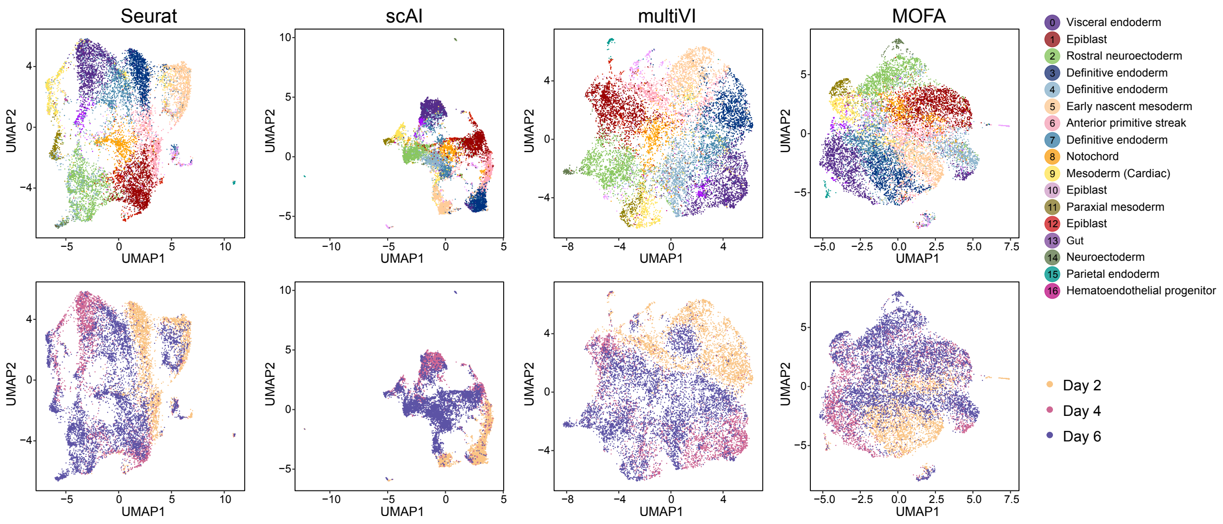


Supplementary Figure S8: UMAP visualization of the dataset for Seurat, scAI, multiVI and MOFA, colored by annotated cell types (first row) and sampling days (second row).

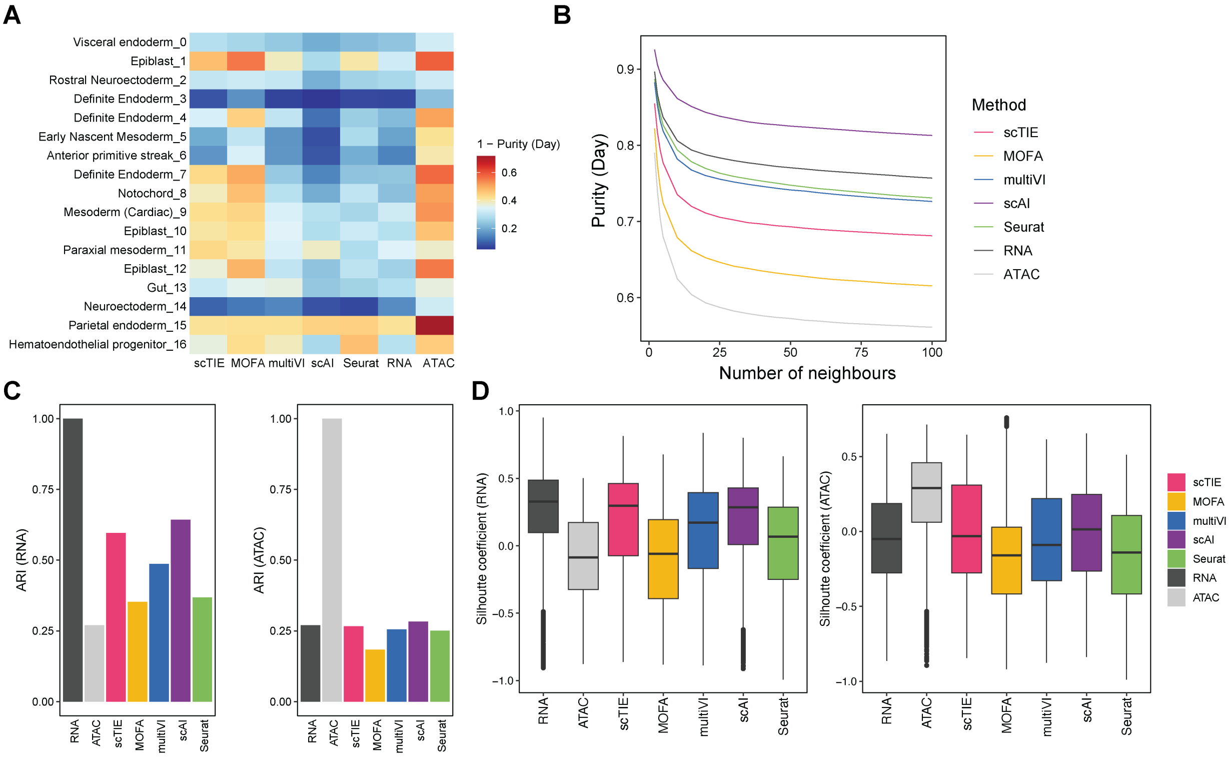


Supplementary Figure S9: (A) 1 - Average purity scores of sampling day for each cell type (Number of neighbors = 50). Row indicates cell types and column indicates methods. Higher values indicate better mixing between days. (B) Average purity scores of sampling day based on different number of neighbors, colored by different methods. Lower values indicate better mixing between days. (C) The bar plots show the ARI values comparing the clustering from different data integration methods with clustering on RNA (left) and ATAC alone (right); higher values indicate better agreement. Note that here RNA and ATAC clustering results are the ground truth for the left panel and right panel respectively, therefore they have ARI equal to 1. (D) Box plots show the silhouette coefficient comparing the clustering from different data integration methods based on distance matrices computed from the RNA (left) and ATAC (right) UMAP coordinates. Higher values indicate better agreement. Note that for the left panel, RNA clustering result has the highest silhouette coefficients because clustering derived from RNA is used as the ground truth; similarly for the right panel.

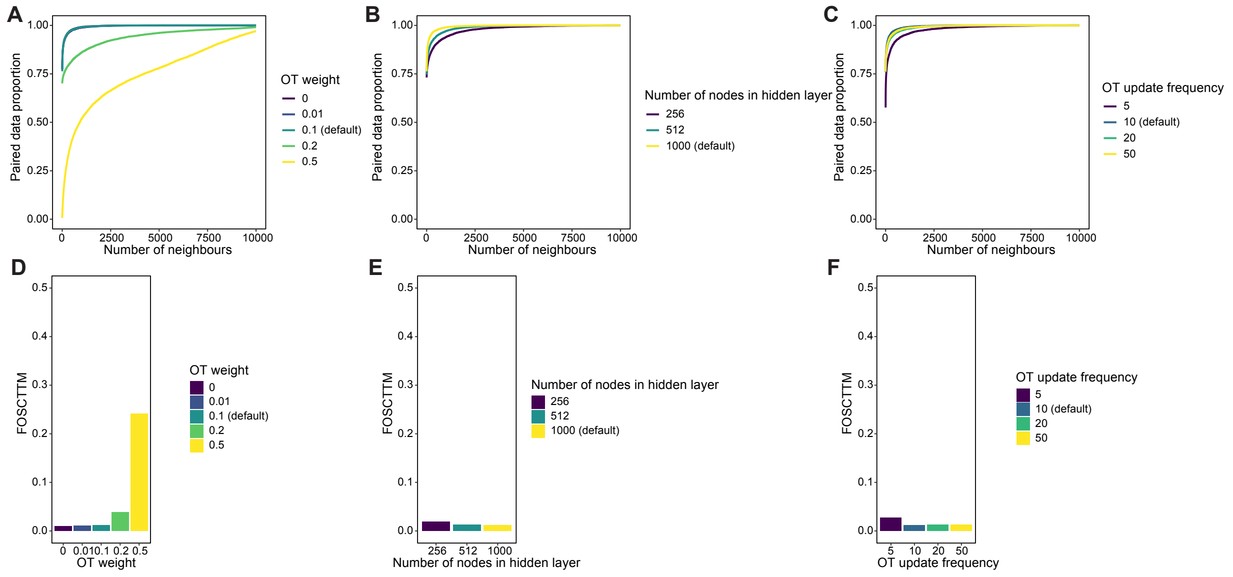


Supplementary Figure S10: Robustness of scTIE with respect to the tuning parameters in modality alignment. (A-C) Proportion of ground truth pairs within certain number of nearest neighbors, with different (A) OT weight; (B) Number of nodes in hidden layer; (C) OT update frequency. (D-F) Barplots of FOSCTTM (fraction of samples closer than the true match) values, with different (D) OT weight; (E) Number of nodes in hidden layer; (F) OT update frequency.

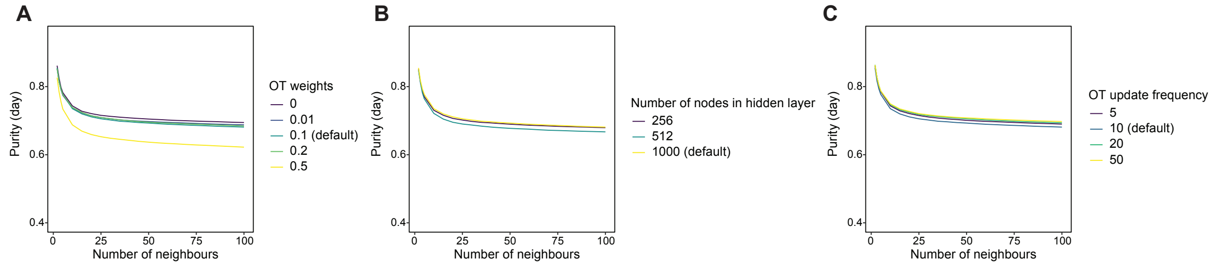


Supplementary Figure S11: Robustness of scTIE with respect to the tuning parameters in time point alignment. Average purity scores of sampling days based on different number of neighbors, with different (A) OT weight; (B) number of nodes in the hidden layer; (C) OT updating frequency.

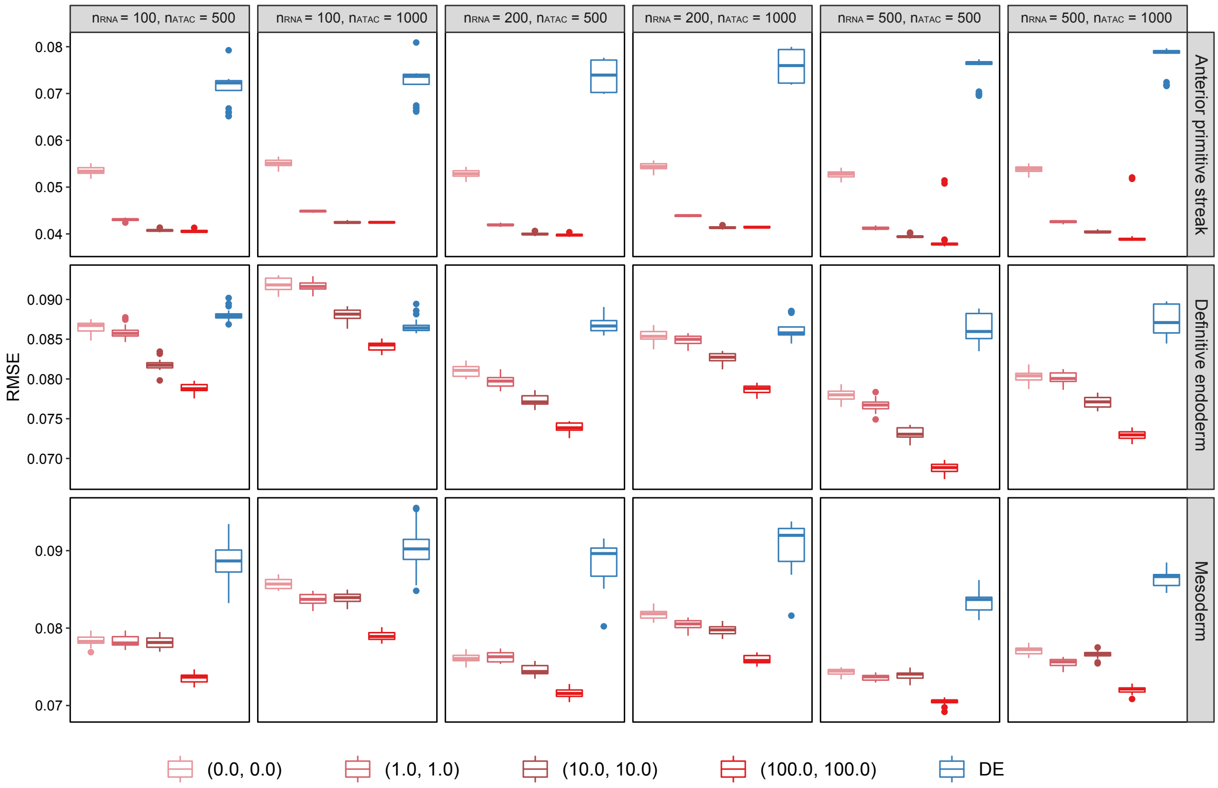


Supplementary Figure S12: Evaluation of transition probability predictions for three different cell fates: anterior primitive streak, definitive endoderm and mesoderm, comparing (1) different number of genes/peaks; (2) different L1 regularization weights used in the prediction task.

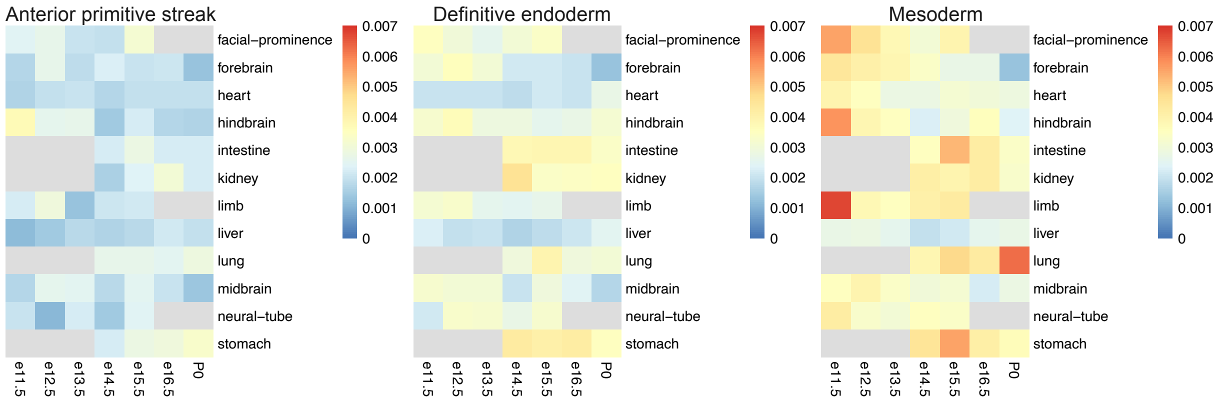


Supplementary Figure S13: Similarity of top DA regions with enhancers of 12 tissues at seven developmental stages from known enhancer databases.

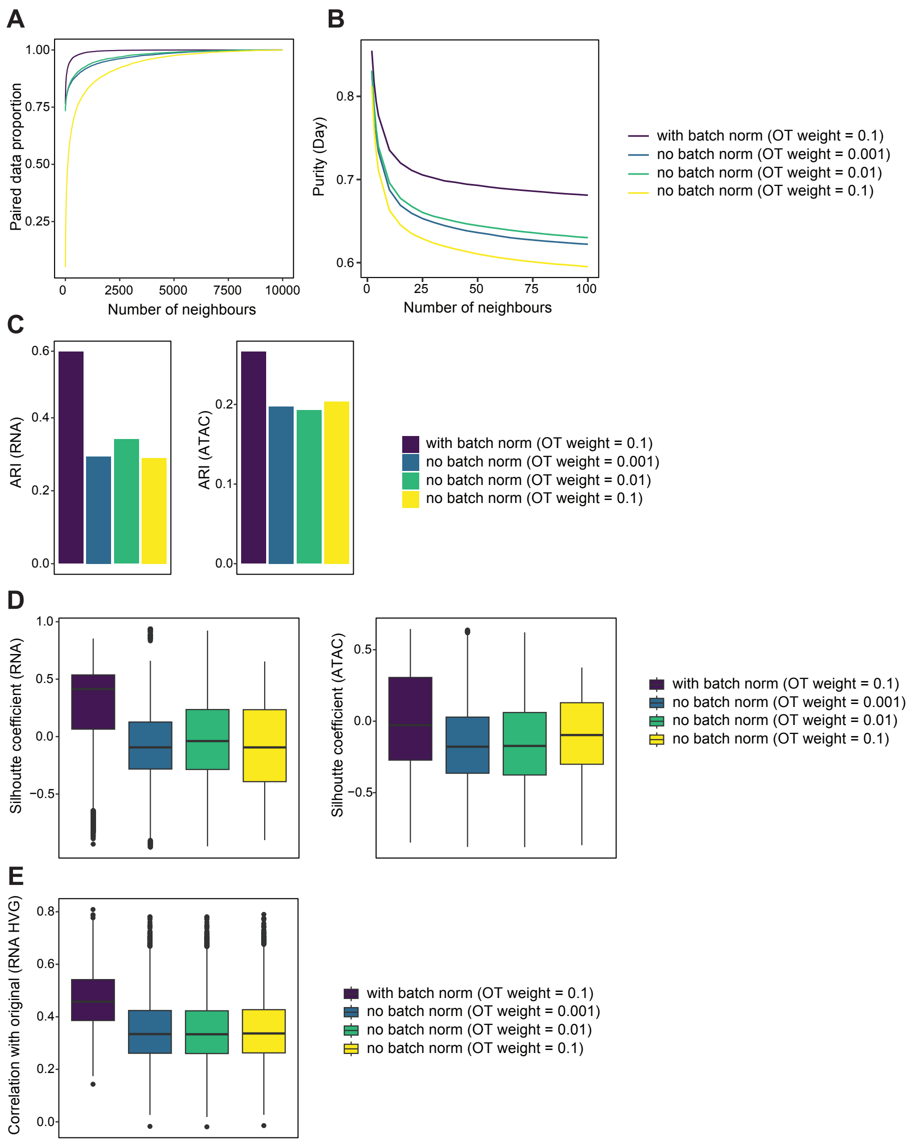


Supplementary Figure S14: scTIE performance comparison with and without the coupled batch norm layers in RNA: (A) Proportion of ground truth pairs within a given number of nearest neighbors; (B) Average purity scores of sampling days based on different numbers of nearest neighbors; (C) The bar plots show the ARI values comparing the clustering from different settings of scTIE with the clustering on RNA (left) and ATAC alone (right); higher values indicate better agreement. (D) Box plots show the silhouette coefficients comparing the clustering from different settings of scTIE based on distance matrices computed from the RNA (left) and ATAC (right) UMAP coordinates. Higher values indicate better agreement. (E) Correlation of scTIE reconstructed RNA expression with the original RNA expression of highly variable genes (HVG).

| | Gradient ranking | DE ranking | DE adj p-value |
|---|---|---|---|
| Sox17 | 126.00 | 1252.00 | 0.46 |
| Smad3 | 260.00 | 2074.00 | 0.81 |
| Zic3 | 150.00 | 1317.00 | 0.49 |
| Twist1 | 441.00 | 184.00 | 0.00 |
| Nfat5 | 368.00 | 2197.00 | 0.84 |
| Hhex | 393.00 | 2650.00 | 0.98 |

Table S1: Comparison of the gradient rankings, DE rankings and adjusted p-values under DE for key TFs in mesoderm lineage.
